# Docking interactions determine substrate specificity of members of a widespread family of protein phosphatases

**DOI:** 10.1101/2024.05.03.592459

**Authors:** Suhaily Caban-Penix, Kristin Ho, Wendy Yang, Rishika Baral, Niels Bradshaw

## Abstract

How protein phosphatases achieve specificity for their substrates is a major outstanding question. PPM family serine/threonine phosphatases are widespread in bacteria and eukaryotes, where they dephosphorylate target proteins with a high degree of specificity. In bacteria, PPM phosphatases control diverse transcriptional responses by dephosphorylating anti-anti-sigma factors of the STAS domain family, exemplified by *B. subtilis* phosphatases SpoIIE, which controls cell-fate during endospore formation, and RsbU, which initiates the General Stress Response. Using a combination of forward genetics, biochemical reconstitution, and AlphaFold2 structure prediction, we identified a conserved, tripartite substrate docking interface comprised of three variable loops on the surface of the PPM phosphatase domain that recognize the three-dimensional structure of the substrate protein. Non-conserved amino acids in these loops facilitate the accommodation of the cognate substrate and prevent dephosphorylation of the non-cognate substrate. Together, single-amino acid substitutions in these three elements cause an over five-hundred fold change in specificity. Our data additionally suggest that substrate-docking interactions regulate phosphatase specificity through a conserved allosteric switch element that controls the catalytic efficiency of the phosphatase by positioning the metal cofactor and substrate. We hypothesize that this is a generalizable mechanistic model for PPM family phosphatase substrate specificity. Importantly, the substrate docking interface with the phosphatase is only partially overlapping with the much more extensive interface with the upstream kinase, suggesting the possibility that kinase and phosphatase specificity evolved independently.

## Introduction

Signaling by reversible phosphorylation requires that opposing kinases and phosphatases have exquisite specificity for their respective substrate proteins (Huse & Kuriyan 2002) (Taylor & Kornev 2011) (Y. Shi 2009). While the mechanisms of kinase specificity, in which sequences surrounding the phosphorylation site dock into a deep active site groove, are well understood (Ubersax & Ferrell 2007), much less is known about how phosphatases discriminate between substrates (Y. Shi 2009). Here, we address the mechanism of how phosphatases achieve substrate specificity with two bacterial serine/threonine phosphatases of the PPM family that must discriminate between their respective substrate proteins in a biological context.

Precisely regulated serine/threonine phosphatases of the PPM family are widespread regulators of bacterial transcriptional responses (Kerk et al. 2015) (Baral et al. 2024). Many organisms have multiple phosphatases that must discriminate between related substrate proteins to maintain signaling fidelity, but the molecular mechanisms of substrate recognition and specificity are not understood (Zhang & L. Shi 2004; Baral et al. 2024; Ho & Bradshaw 2021; Bradshaw et al. 2017). A particular challenge to determining how PPM phosphatases achieve specificity is that the active site is accessible on the solvent-exposed surface. Here we determine the mechanism of specificity for two phosphatases from *B. subtilis*. SpoIIE dephosphorylates SpoIIAA to specify cell fate during endospore formation by activating σ^F^ (Stragier & Losick 1996; Duncan et al.1995), and RsbU dephosphorylates RsbV to initiate the general stress response by activating σ^B^ (Yang et al. 1996) (Fig. 1A). SpoIIAA and RsbV are paralogues that share the STAS domain fold and share 29 percent sequence identity (Seavers et al. 2001) (Kovacs et al. 1998; Pathak et al. 2020; Sharma et al. 2011).

**Figure 1:**
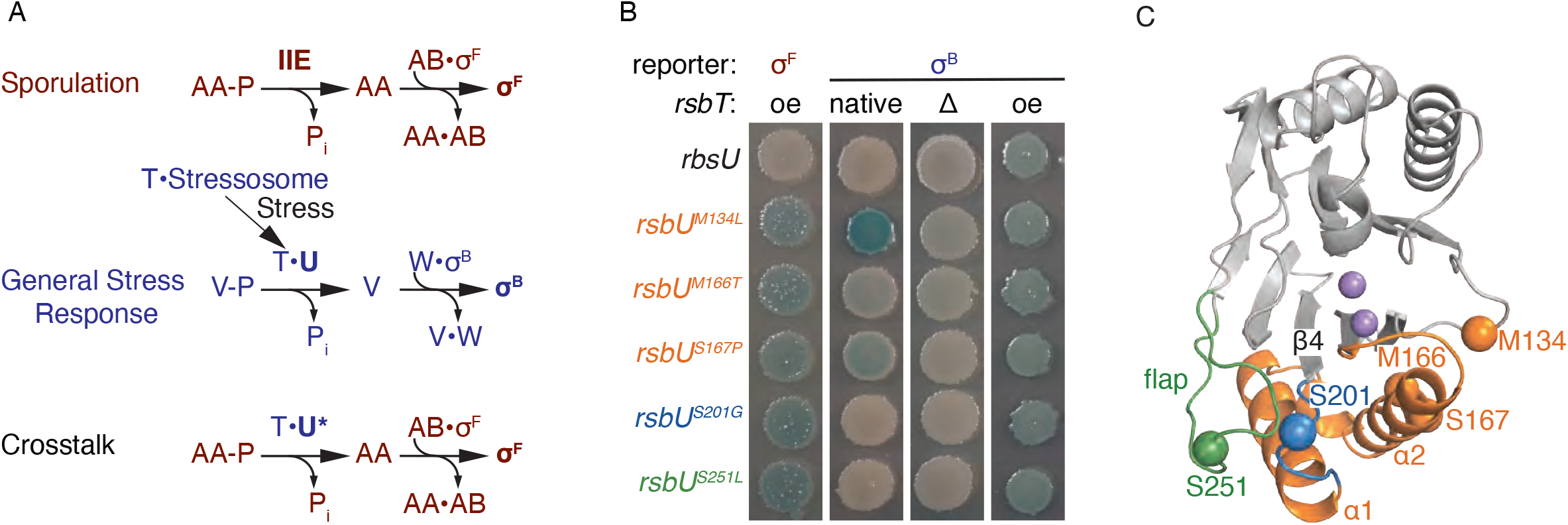
Isolated crosstalk mutants fall into two phenotypic classes. **A**. A depiction of the sporulation (red) and general stress response pathways (blue), along with the premise for our genetic screen in looking for crosstalk mutants. PPM phosphatases, SpoIIE (red), and RsbU (blue) dephosphorylate paralogous substrates SpoIIAA (red) and RsbV (blue), respectively. Activation of RsbU is dependent on RsbT. Dephosphorylation of SpoIIAA and RsbV activates a partner-switching mechanism, where the unphosphorylated substrates SpoIIAA and RsbV bind to the kinases SpoIIAB (red) and RsbW (blue). The binding of the substrates to the kinases releases the sigma factors σ^F^ (red) and σ^B^ (blue). A screen was developed to isolate RsbU crosstalk mutants that could activate σ^F^ in a strain that was deleted for *spoIIE*. **B**. RsbU variants with amino acid substitutions (M134L, M166T, S167P, S201G, and S251L) isolated in the genetic screen activate σ^F^ and retain their ability to activate σ^B^ in *B. subtilis*. Reporter strains with *lacZ* under the control of either the σ^F^ (left) or σ^B^ (right) promoter were plated on indicator plates containing x-gal and IPTG. The first σ^B^ reporter strain has *rsbT* on the chromosome (left), the second *rsbT* is deleted (middle) and the last *rsbT* is overexpressed (oe) from plasmid pHB201. Plates were imaged after 24 hours of growth at 37°C. **C**. AlphaFold2 structure of RsbU phosphatase domain. Residues M166 and S167 are found in the switch (orange), which includes the α1 and α2 helixes. M134 (orange) is adjacent to the switch. S201 (blue) is in the loop between α1(orange) helix and β4 (gray). S251 is located in the flap (green). The two metals sit at the catalytic center (purple).

Three features of these systems make them ideal for understanding molecular mechanisms of phosphatase specificity. First, crosstalk between these pathways is highly detrimental; activation of σ^B^ by SpoIIE blocks sporulation (Rothstein et al. 2017) and activation of σ^F^ by RsbU causes lethality (Fig. S1). Second, each phosphatase acts on a single phospho-serine on a single substrate protein, simplifying the analysis of changes in specificity in both cellular and biochemical contexts(Najafi et al. 1995; Yang et al. 1996; Carniol, Kim, et al. 2004). Third, we have previously biochemically reconstituted the specificity of both phosphatases and have found them to be highly specific (SpoIIE has an approximately 4000-fold greater k_cat_/K_M_ for SpoIIAA and RsbU has approximately an 400-fold greater k_cat_/K_M_ for RsbV) (Ho & Bradshaw 2021).

An important consideration for phosphatase specificity is that every substrate of a protein-phosphatase is shared with an opposing kinase. An unanswered question is whether the kinases and phosphatases recognize the same or different features of their shared substrate proteins. Whether the same features of the substrate proteins are recognized by both enzymes determines how changes in substrate sequence impact signaling and constrain the pathways available for evolving new signaling functions. SpoIIAA and RsbV are each phosphorylated by a cognate kinase/anti-sigma factor (SpoIIAB and RsbW respectively) that undergoes a partner-switch to release the sigma factor when the substrate is dephosphorylated (Garsin et al. 1998; Yang et al. 1996) (Fig. 1A). X-ray crystal structures revealed extensive interfaces surrounding the phosphorylation sites for both SpoIIAA/SpoIIAB and RsbV/RsbW complexes, enabling direct comparison of how the phosphatases and kinases recognize their substrates (Pathak et al. 2020; Masuda et al. 2004).

PPM family phosphatases use two divalent cations in their active sites to deprotonate a water that is the nucleophile for attack of the phospho-serine (Das et al. 1996; Y. Shi 2009). Our structural and biochemical studies revealed that the activity of SpoIIE and RsbU is controlled by the conformational change of an α-helical switch element at the base of the phosphatase domain (α1 and α2) that coordinates the metal cofactor (Baral et al. 2024; Ho & Bradshaw 2021; Bradshaw et al. 2017). Subsequent genetic and biochemical experiments implicated this element in substrate specificity, but the molecular mechanistic basis for this was not known (Ho & Bradshaw 2021). One clue is that a variable insertion region termed the “flap” that has been implicated in other systems packs against the switch element, suggesting that substrate docking could be transmitted through these contacts (Pullen et al. 2004; Bradshaw et al. 2017; Gilmartin et al. 2014; Waschbüsch et al. 2021; Yamaguchi et al. 2005). However, the binding interface of the substrate protein and phosphatase had not been identified.

Using a combination of genetics, biochemical reconstitution, and AlphaFold2 structure prediction, we have discovered the molecular basis for how SpoIIE and RsbU specifically recognize their respective substrate proteins. They use a conserved tri-partite binding site where the folded protein substrate engages with three variable loops of the phosphatase domain that dock against the switch element to position the substrate and form the catalytic center. We hypothesize that this is a broadly generalizable mechanism by which PPM family phosphatases engage with and achieve specificity for their substrates.

## Results

### Isolation of RsbU crosstalk mutants

To identify the features of RsbU that determine substrate specificity, we designed a genetic screen to isolate crosstalk variants of RsbU that dephosphorylate the off-pathway substrate, SpoIIAA, and activate σ^F^ (Fig. 1A). We introduced plasmids with PCR mutagenized *rsbT* and *rsbU* genes to a σ^F^ reporter strain lacking the σ^B^ operon (including *rsbT* and *rsbU*), and *spoIIE*, the phosphatase responsible for activating σ^F^. We then screened for plasmids that cause activation of σ^F^ under sporulation conditions when *rsbT* and *rsbU* are expressed. Performing the screen under sporulation conditions was essential because uncompartmentalized activation of σ^F^ is lethal (Fig. S1). Because expression of the SpoIIA operon, which includes σ^F^, occurs only in a sub-population of cells during sporulation, this allows isolation of cells carrying plasmids that drive improper activation of σ^F^. After an initial round of screening did not yield any crosstalk mutations, we increased the sensitivity of our screen by decreasing the activity of the SpoIIAB kinase (using a strain with *spoIIAB^R105C^*) (Carniol, Eichenberger, et al. 2004). We identified five *rsbU* mutants that caused robust σ^F^ activation from this screen (M134L, M166T, S167P, S201G, and S251L), but did not isolate mutations in *rsbT* (Fig 1B). All amino acid substitutions mapped to the phosphatase domain of RsbU. While two *rsbU* variants were only isolated from one pool of mutagenized plasmids, the others were picked up from two or more independently generated plasmid pools, suggesting that the screen was near saturation.

### Crosstalk mutants fall into two phenotypic classes

There are three possible models for how *rsbU* crosstalk mutations cause σ^F^ activation in our screen. First, crosstalk mutations could swap RsbU specificity, increasing activity towards SpoIIAA and decreasing activity towards RsbV. Second, crosstalk mutations could cause indiscriminate dephosphorylation of both SpoIIAA and RsbV. Third, crosstalk mutations could hyperactivate RsbU, leading to SpoIIAA dephosphorylation without a change in specificity. To qualitatively distinguish between these models, we transformed plasmids containing rebuilt versions of the *rsbU* crosstalk mutants into additional reporter strains.

First, to determine whether any RsbU variants lost activity towards RsbV, we overexpressed the *rsbU* variants in a σ^B^ reporter strain deleted for *rsbTU* on the chromosome. All five *rsbU* mutants robustly activated σ^B^, similar to wild-type *rsbU*, indicating that they retain activity towards RsbV and are at least somewhat promiscuous (Fig 1B).

Second, to determine whether any of the RsbU mutants had increased activity towards RsbV, we rebuilt the *rsbU* mutations in a plasmid that did not contain *rsbT* and transformed these plasmids into σ^B^ reporter strains. *rsbU^M134L^*, *rsbU^M166T^*, and *rsbU^S167P^* activated σ^B^ in a strain background where *rsbT* was present on the chromosome, while none of the *rsbU* variants activated σ^B^ in a strain deleted for *rsbT* (Fig 1B). From this we conclude that the *rsbU* variants fall into two classes: M134L, M166T, and S167P are hyperactivating, while S201G and S251L are not.

### A structural model for phosphatase/substrate interaction

To further characterize the RsbU crosstalk variants, we mapped their locations onto the RsbU phosphatase domain from an AlphaFold2 model that we generated of dimeric RsbU (Baral et al. 2024) (Fig. 1C). The mutations cluster in two regions:

M134, M166 and S167 form a cluster, buried in the core of the phosphatase domain around the α1 helix (Fig. 1C). The α1 and α2 helices control phosphatase activation and substrate recognition in the paralogous phosphatase, SpoIIE, suggesting that this mechanism is conserved with RsbU (Baral et al. 2024; Ho & Bradshaw 2021; Bradshaw et al. 2017). Interestingly, we previously isolated substitutions at M166 in a screen to identify RsbT-independent variants of RsbU (Ho & Bradshaw 2021), and in a suppressor screen to restore activity to an RsbU variant that has reduced binding to RsbT (Baral et al. 2024). To determine whether the identity of the amino acid substitution at M166 differentially impacted σ^F^ and σ^B^ activity, we generated an allelic series replacing M166 with amino acids of varied size and hydrophobicity. We found that patterns of σ^F^ and σ^B^ activation generally paralleled each other, suggesting that hyperactivity and promiscuity are related (Fig. S2A).

S201 and S251 are located on the opposite side of the active site from M166and are predicted to be solvent-exposed, suggesting that they could make direct contact with RsbV (Fig. 1C). S201 is in the loop between α2 and β4, while S251 is located in a variable element of the phosphatase domain (referred to as the “flap”) that has been implicated in substrate recognition in other PPM phosphatases(Pullen et al. 2004; Bradshaw et al. 2017; Gilmartin et al. 2014; Waschbüsch et al. 2021; Yamaguchi et al. 2005). This conclusion is supported by our AlphaFold2 model of a heterotetrameric RsbU/V complex that places S201 and S251 at the RsbU/V interface (Fig 2A). The phosphorylation site, S56, is modeled near the catalytic center of RsbU, additionally supporting the validity of the model. RsbV is predicted to exclusively contact the phosphatase domain of RsbU and would not contact RsbT in the hetero-heptameric RsbT/U/V signaling complex. Thus, we conclude that S201 and S251 are likely to be directly involved in substrate recognition.

**Figure 2:**
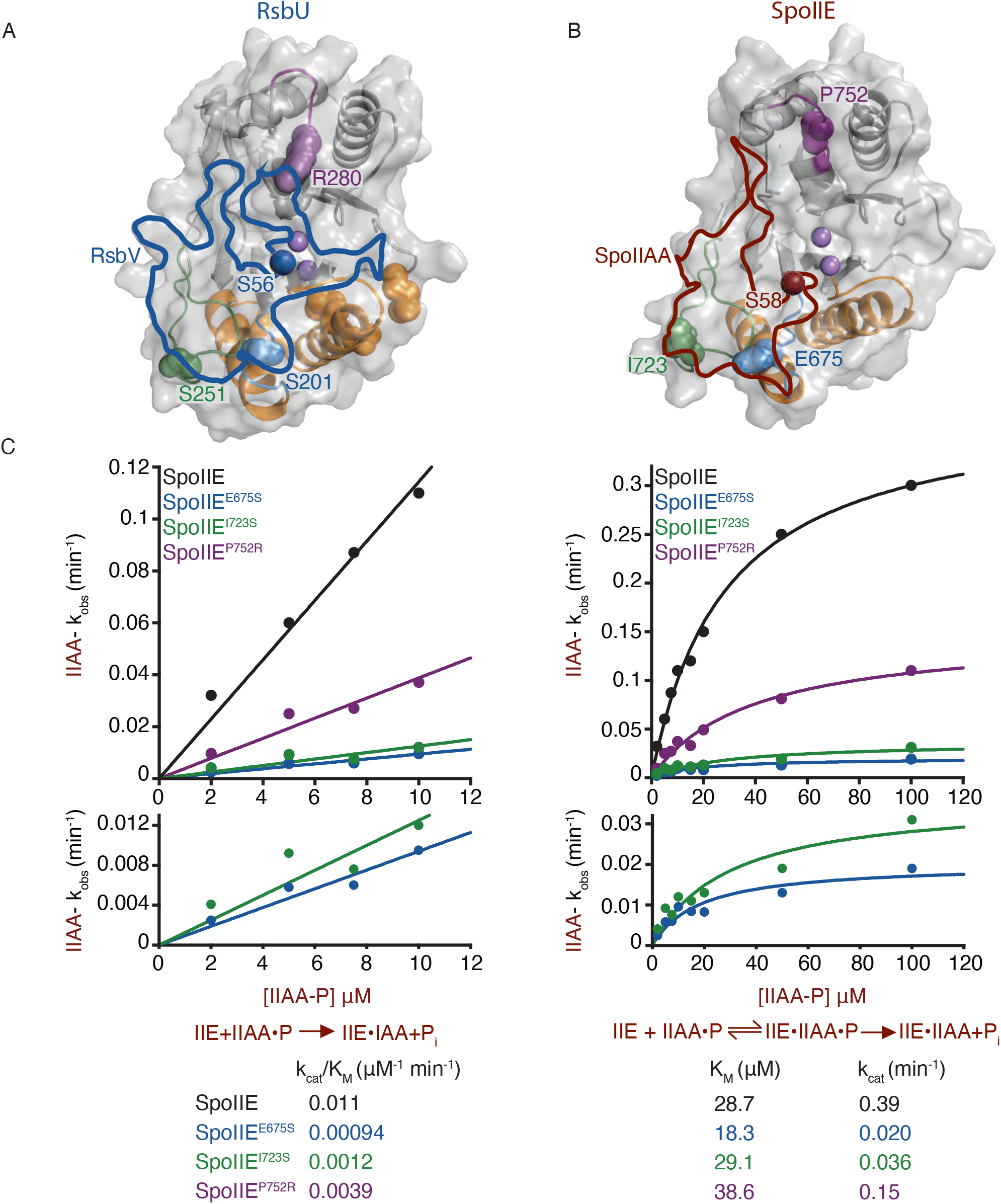
Conservation at the phosphatase/substrate interface positions SpoIIAA for efficient catalysis. **A.** Alphafold2 structure of RsbU phosphatase domain. The position of RsbV (blue) at the RsbU/RsbV interface is outlined based on a 1.4 Å probe radius with a transparent surface model shown. RsbV phosphorylation site, S56 (red) sites near the catalytic center where the two metals are coordinated (light purple). Residue R280 is located in the α3/4 loop. **B.** Alphafold2 structure of SpoIIE phosphatase domain. The position of SpoIIAA at the SpoIIE/SpoIIAA interface is outlined (red) with a 1.4 Å probe radius. SpoIIAA phosphorylation site, S58 (red), is located near the catalytic center in proximity to the two metal cofactors (light purple). Residues E675S (blue), I723 (green), and P752 (purple) are located within the α1/β4 loop, flap, and α3/4 loop, respectively. **C**. Graphs showing the rate of SpoIIE (black), SpoIIE^E675S^ (blue), SpoIIE^I723S^ (green), and SpoIIE^P752R^ (purple) dephosphorylation as a function of SpoIIAA-P concentration. The plot on the left displays the k_cat_/K_M_ values of SpoIIAA-P dephosphorylation by SpoIIE (black), SpoIIE^E675S^ (blue), SpoIIE^I723S^ (green), and SpoIIE^P752R^ (purple) with concentrations of SpoIIAA-P below the K_M_ fit to the linear equation (in KaleidaGraph) (k_cat_/K_M_)*[SpoIIAA-P]. The k_cat_/K_M_ were SpoIIE 0.011± 0.00045 µM^-1^ min^-1^, SpoIIE^E675S^ 0.001± 7.0e^-5^ µM^-1^ min^-1^, SpoIIE^I723S^ 0.001± 0.00016 µM^-1^ min^-1^, and SpoIIE^P752R^ 0.0039± 0.00028 µM^-1^ min^-1^. The plot on the right was fit to the Michaelis-Menten equation (in KaleidaGraph) k_cat_*[SpoIIAA-P]/(K_M_ + [SpoIIAA-P]). The k_cat_ for each enzyme were SpoIIE 0.39±0.017 min^-1^, SpoIIE^E675S^ 0.020± 0.0026 min^-1^, SpoIIE^I723S^ 0.036± 0.0058 min^-1^, and SpoIIE^P752R^ 0.15± 0.014 min^-1^. The K_M_ measured for each enzyme were SpoIIE 28.7± 2.8 µM, SpoIIE^E675S^ 18.3± 6.0 µM, SpoIIE^I723S^ 29.1± 10.5 µM, and SpoIIE^P752R^ 38.6± 7.9 µM. The error is the error of the fit. Reactions were multiple turnover reactions with varying concentrations of SpoIIAA-P, 0.1 µM SpoIIE, 10 mM MgCl_2_, and 0.1 µM SpoIIAA-P^32^. Below each graph is a summary of the reaction. On the left are the kinetic parameters for the k_cat_/K_M_ reaction scheme, P_i_ indicating product, and kobs values below. On the right, the kinetic scheme is used to summarize the parameters and values for k_cat_ and K_M_ values.

### Conservation of the phosphatase/substrate interface

Next, to assess whether the RsbU/V interface is shared with SpoIIE and SpoIIAA, we generated a similar AlphaFold2 model of a heterotetrameric SpoIIE/AA complex based on the dimeric structure of SpoIIE that we determined previously (Bradshaw et al. 2017) (Fig. 2B). The model places SpoIIAA in a very similar position relative to the SpoIIE phosphatase domain as we observed in the RsbU/V complex (Fig. 2A). Importantly, SpoIIE residues corresponding to crosstalk variants RsbU^S201^ (SpoIIE^E675^) and RsbU^S251^ (SpoIIE^I723^) are buried in the interface. Additionally, both of these residues stand out as being variable elements of the contact interface, which is otherwise relatively conserved (Fig. S2B). One notable difference between the models is that SpoIIAA makes contacts, distant from the phosphorylation site, with the regulatory domain of SpoIIE (Fig. S2B). These contacts provide an explanation for why mutation of an amino acid in this interface (glutamine 73 to alanine, SpoIIAA^Q73A^) causes hyperactivation of σ^F^ (Carniol, Eichenberger, et al. 2004), and further supports the validity of the structural model.

### Flap and switch loops position SpoIIAA for dephosphorylation

Next, we biochemically assayed how the contacts identified by our genetic screen and structural models determine substrate specificity. We used SpoIIE for these studies because we have more extensively studied SpoIIE specificity compared to RsbU, and the K_M_ of RsbU is below 1µM for both SpoIIAA and RsbV, making measurement of k_cat_/K_M_ more challenging. We generated variants of the phosphatase domain of SpoIIE (SpoIIE^590-827^, which we previously found is sufficient to recapitulate substrate specificity (Ho & Bradshaw 2021)) that were substituted for the corresponding amino acid of RsbU at positions E675 (serine) and I723 (serine). Other than these non-conserved interface residues that were genetically identified, we selected one additional non-conserved interface residue to mutate based on analysis of conservation of the phosphatase/substrate interface predictions, SpoIIE^P752R^ (Fig. 2A-B, Fig. S2C). P752 repositions a variable loop above the phosphatase active sites (RsbU^279-285^/SpoIIE^751-754^) that forms a contact with RsbV in the RsbU/RsbV complex model. We therefore hypothesized that the SpoIIE^P752R^ mutation might favor recognition of RsbV.

Using an assay that monitors dephosphorylation of ^32^P-SpoIIAA, we found that SpoIIE^E675S^ (k_cat_^SpoIIAA^/K_M_^SpoIIAA^ 0.001 µM^-1^min^-1^) and SpoIIE^I723S^ (k_cat_^SpoIIAA^/K_M_^SpoIIAA^ 0.001µM^-1^min^-1^) had ten-fold reductions in k_cat_^SpoIIAA^/K_M_^SpoIIAA^ compared to SpoIIE (0.011 µM^-1^min^-1^) (Fig. 2C). Extending these data to near saturating concentrations of SpoIIAA revealed that the primary effect of the substitutions was on the k_cat_^SpoIIAA^ and that there was no significant change in K_M_^SpoIIAA^ (Fig. 2C). Consistent with the defects in phosphatase activity being substrate specific, we observed no change in the activity towards the generic small-molecule substrate, p-nitrophenyl phosphate (Fig. S3). Thus, we conclude that variable positions in the flap and switch regions mediate substrate contacts important for achieving maximal catalytic efficiency for the cognate substrate.

The specificity and activity of SpoIIE and RsbU is additionally determined by recruitment of metal cofactor (Ho & Bradshaw 2021; Bradshaw et al. 2017). Thus, we held the substrate concentration constant and measured SpoIIE activity as a function of metal concentration (Fig. S4). None of the variants had a significant change in metal concentration dependence of activity, establishing that they do not impact this step of the reaction. We additionally explored the metal cofactor preference of the SpoIIE variants. Our initial assays were performed with magnesium, which we presume is the physiologically relevant metal. However, the phosphatase domain construct of SpoIIE is more active with manganese than magnesium, so we assayed SpoIIE^E675S^, one of the variants with the largest effects with manganese as the metal cofactor. In this case, we observed only a two-fold decrease in activity, suggesting that the identity of the metal cofactor influences the effect of specificity determinants (Fig. S5). Together, we conclude that specificity determinants in the flap and switch regions support efficient catalysis of cognate substrate once bound in a manner that depends on the identity of the metal cofactor.

### The switch and α3/4 loop discriminate against RsbV

Specificity is determined by the relative k_cat_/K_M_ for two competing substrates. Thus, to determine the contributions of the switch loop (SpoIIE-E675), flap (SpoIIE-I723), and α3/4 loop (SpoIIE-P752) to specificity, we measured hydrolysis of ^32^P-RsbV by SpoIIE variants. We conducted these assays with manganese as the metal cofactor because the rate of hydrolysis with magnesium was too slow to accurately measure. We found that SpoIIE^P752R^ (k_cat_^RsbV^/K_M_^RsbV^ 0.0035 µM^-1^min^-1^) was 17-fold more active towards RsbV-P than SpoIIE (k_cat_^RsbV^/K_M_^RsbV^ 0.00045 µM^-1^min^-1^), while SpoIIE^E675S^ (k_cat_^RsbV^/K_M_^RsbV^ 0.00058 µM^-1^min^-1^) and SpoIIE^I723S^ (k_cat_^RsbV^/K_M_^RsbV^ 0.00028 µM^-1^min^-1^) did not significantly change activity (Fig. 3B). Extending the data to higher concentrations of SpoIIE demonstrated that the P752R substitution primarily increases the k_cat_^RsbV^ without changing the K_M_^RsbV^ (although we were not able to saturate the reaction due to insolubility of SpoIIE^P752R^ at high concentrations). We additionally observed some increase in k_cat_^RsbV^ for SpoIIE^E675S^ and SpoIIE^I723S^, but these effects were offset by increases in K_M_^RsbV^. Of note, we postulate that the magnitude of the effect could be an underestimate because our experiments with SpoIIAA suggest that SpoIIE is more promiscuous when manganese is used as the metal cofactor compared to magnesium. We conclude that the α3/4 loop is important for substrate discrimination and that the P752R substitution allows SpoIIE to accommodate the non-cognate substrate, RsbV-P, in a manner more favorable for catalysis (Fig. 3A).

**Figure 3:**
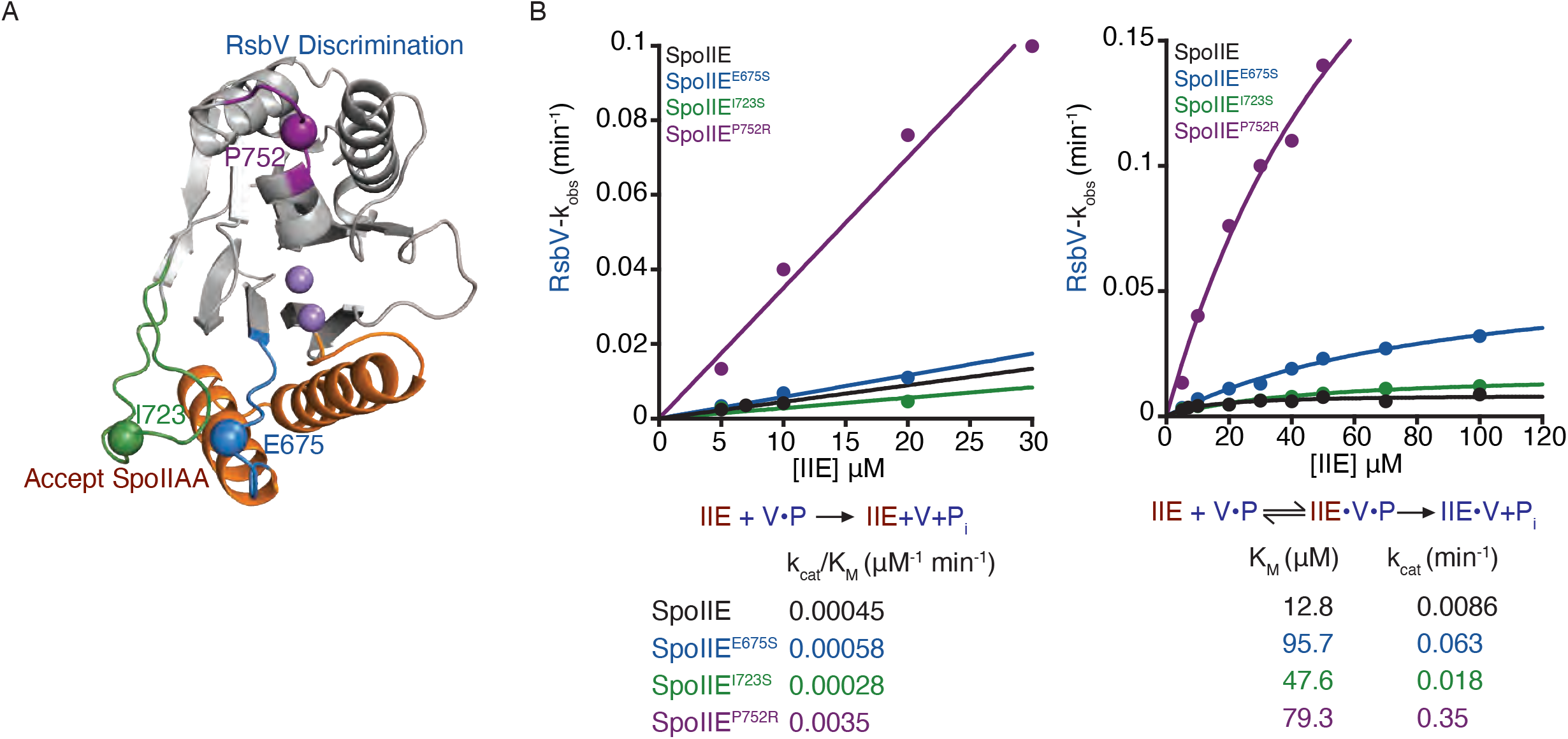
The α3/4 loop contributes to specificity by discriminating against RsbV. **A**. AlphaFold2 structure of SpoIIE phosphatase domain. The SpoIIE phosphatase domain shows the positions of residues I723 (green), E675S (blue), P752 (purple), and metal cofactors (light purple) depicted on the structure. The rejection of RsbV (blue) by the residue P752 and acceptance of SpoIIAA (red) by residues I723S and E675 are noted on the structure. **B.** Graphs showing the rate of SpoIIE, SpoIIE^E675S^, SpoIIE^I723S^, and SpoIIE^P752R^ dephosphorylation of RsbV-P. The plot on the *right* displays the k_cat_/K_M_ values of RsbV-P dephosphorylation by SpoIIE (black), SpoIIE^E675S^ (blue), SpoIIE^I723S^ (green), and SpoIIE^P752R^ (purple) with concentrations of RsbV-P below the K_M_ fit to a linear equation (in KaleidaGraph) (k_cat_/K_M_)*[RsbV-P]. The k_cat_/K_M_ were SpoIIE 0.00045± 2.9e-5 µM^-1^ min^-1^, SpoIIE^E675S^ 0.00058± 3.8e^-5^ µM^-1^ min^-1^, SpoIIE^I723S^ 0.00028± 6.5e-5 µM^-1^ min^-1^, and SpoIIE^P752R^ 0.0035± 0.00028 µM^-1^ min^-1^.The plot on the *left* was fit to the Michaelis-Menten equation (in KaleidaGraph) kcat*[RsbV-P]/(K_M_ + [RsbV-P]). The k_cat_ for each enzyme were SpoIIE 0.0086± 0.00080 min^-1^, SpoIIE^E675S^ 0.063± 0.0071 min^-1^, SpoIIE^I723S^ 0.018± 0.0021 min^-1^, and SpoIIE^P752R^ 0.15± 0.096 min^-1^. The K_M_ measured for each enzyme were SpoIIE 12.8± 4 µM, SpoIIE^E675S^ 95.7± 17.5 µM, SpoIIE^I723S^ 47.6± 12.0 µM, and SpoIIE^P752R^ 79.3± 31.6 µM. The error is the error of the fit. Reactions were single turnover reactions with varying concentrations of SpoIIE, 0.15 µM RsbV-P, 10 mM, and MnCl_2_. Below each graph is a summary of the reaction. On the left are the kinetic parameters for the k_cat_/K_M_ reaction scheme, P_i_ indicating product, and kobs values below. On the right, the kinetic scheme is used to summarize the parameters and values for k_cat_ and K_M_ values.

### Combinatorial control of substrate discrimination

Together, we identified three specificity determinants in the phosphatase domain of SpoIIE that recognize SpoIIAA (the switch-loop and flap) and reject RsbV (the α3/4 loop). To determine how these features work together, we generated a triple-mutant variant that combines all three substitutions. For this variant, we observed a combinatorial effect, with a hundred-fold reduction in k_cat_^SpoIIAA^/K_M_^SpoIIAA^ (Fig. 4A, B), and a roughly five-fold increase in k_cat_^RsbV^/K_M_^RsbV^ (Fig. 4C). Although this is not a direct comparison (because different metal cofactors were used with SpoIIAA and RsbV substrates), these changes represent a substantial reduction in the specificity of SpoIIE, while boosting the activity towards the non-cognate substrate. Sequence alignments of diverse bacterial phosphatases reveal that these three loop elements are variable in sequence and length, supporting a model that they are shared determinants of substrate specificity.

**Figure 4:**
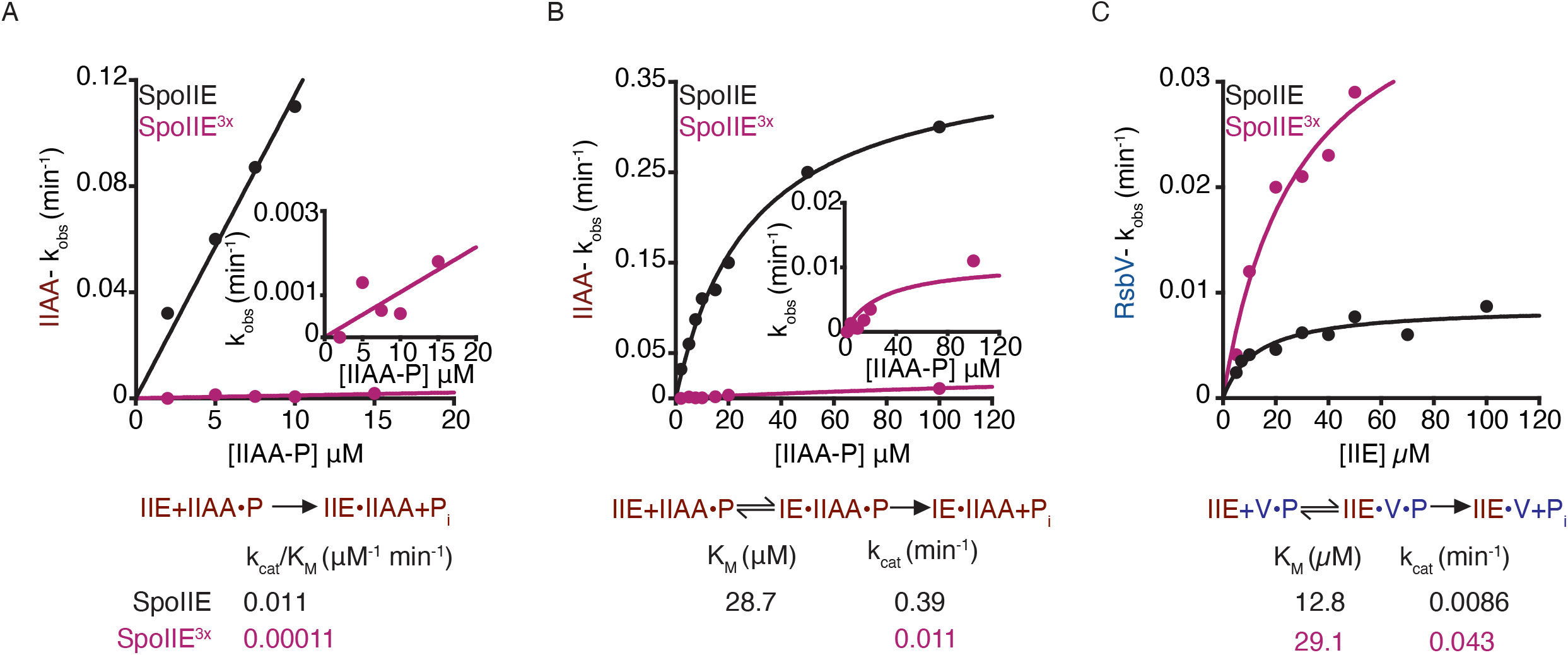
Combination of all three amino acid substitutions substantially reduces substrate specificity. All plots show the rates of SpoIIAA-P dephosphorylation by SpoIIE and SpoIIE^3x^ as a function of SpoIIAA-P concentration. **A.** The k_cat_/K_M_ values SpoIIAA-P dephosphorylation by SpoIIE (black) and SpoIIE^3x^ (pink) with concentrations of SpoIIAA-P below the K_M_ were fit to the linear equation (in KaleidaGraph) (k_cat_/K_M_)*[SpoIIAA-P]. The k_cat_/K_M_ were SpoIIE 0.011± 0.00045 µM^-1^ min^-1^, SpoIIE^3x^ 0.00011± 2.4e^-5^ µM^-1^ min^-1^. The error is the error of the fit. Reactions were multiple turnover reactions with varying concentrations of SpoIIAA-P, 0.1 µM SpoIIE, 10 mM MgCl_2_, and 0.1 µM SpoIIAA-P^32^. Below the graph is a summary of the reaction scheme depicting the kinetic parameters for k_cat_/K_M_, P_i_, the product, and k_obs_ values below. **B.** Rate of SpoIIAA-P dephosphorylation by SpoIIE (black) and SpoIIE^3x^ (pink) as a function of SpoIIAA-P concentration. The plot was fit to the Michaelis-Menten equation (in KaleidaGraph) k_cat_*[SpoIIAA-P]/(K_M_ + [SpoIIAA-P]) using a non-linear curve fitting. The k_cat_ measured was SpoIIE 0.39± 0.017 min^-1^ and SpoIIE^3x^ 0.011± 0.0018 min^-1^. The error is the error of the fit. The K_M_ for SpoIIE was 0.39± 2.8 µM. Reactions were multiple turnover reactions with varying concentrations of SpoIIAA-P, 0.1 µM SpoIIE, 10 mM MgCl_2_, and 0.1 µM SpoIIAA-P^32^. Below the graph is a summary of the reaction scheme depicting the kinetic parameters for k_cat_, K_M_, P_i_, the product, and k_obs_ values below. **C.** Rate of RsbV dephosphorylation by SpoIIE (black) and SpoIIE^3x^ (pink) as a function of SpoIIE concentration. The plot was fit to a Michaelis-Menten equation (in KaleidaGraph) k_cat_*[SpoIIAA-P]/(K_M_ + [SpoIIAA-P]) using a non-linear curve fitting. The k_cat_ measured was SpoIIE 0.0086± 0.00080 min^-1^, and SpoIIE^x3^ 0.043± 0.0084 min^-1^. The K_M_ measured was SpoIIE 12.8± 4.0 µM and SpoIIE^x3^ 29.1± 11.8 µM. The error is the error of the fit. Reactions were single turnover reactions with varying concentrations of SpoIIE, 0.15 µM RsbV-P, 10 mM, and MnCl_2_. Below the graph is a summary of the reaction scheme depicting the kinetic parameters for k_cat_, K_M_, P_i_, the product, and k_obs_ values below.

### Substrate complementarity drives phosphatase specificity

To determine how features of the substrate protein interact with specificity determinants in the phosphatase domain, we generated a variant of SpoIIAA with arginine 67 substituted with threonine (the homologous residue of RsbV). We found that SpoIIAA^R67T^ was dephosphorylated roughly twenty-fold slower by SpoIIE (measured under k_cat_/K_M_ conditions) (Fig. 5A). Remarkably, R67 of SpoIIAA is predicted to be in proximity of E675 of SpoIIE in our AlphaFold2 model of the SpoIIAA/SpoIIE complex (Fig. 5B). We therefore assayed dephosphorylation of SpoIIAA^R67T^ by SpoIIE^E675S^ to determine if the substitutions were compensatory. Indeed, we found that the activity of SpoIIE^E675S^ was greater towards SpoIIAA^R67T^ than SpoIIAA (Fig. 5A). Importantly, this was not the case for SpoIIE^I723S^, for which we observed a two-fold reduction in rate regardless of the substrate (Fig. S6). This finding provides further support for the AlphaFold2 model of the SpoIIE/SpoIIAA complex and emphasizes the importance of the loop connecting α2 (part of the switch element) to β4 of the PPM fold for specificity. The position of R67 on SpoIIAA additionally demonstrates that docking interactions involving the three-dimensional structure, distant from the phosphorylation site of the substrate protein, are critical for recognition by the phosphatase.

**Figure 5:**
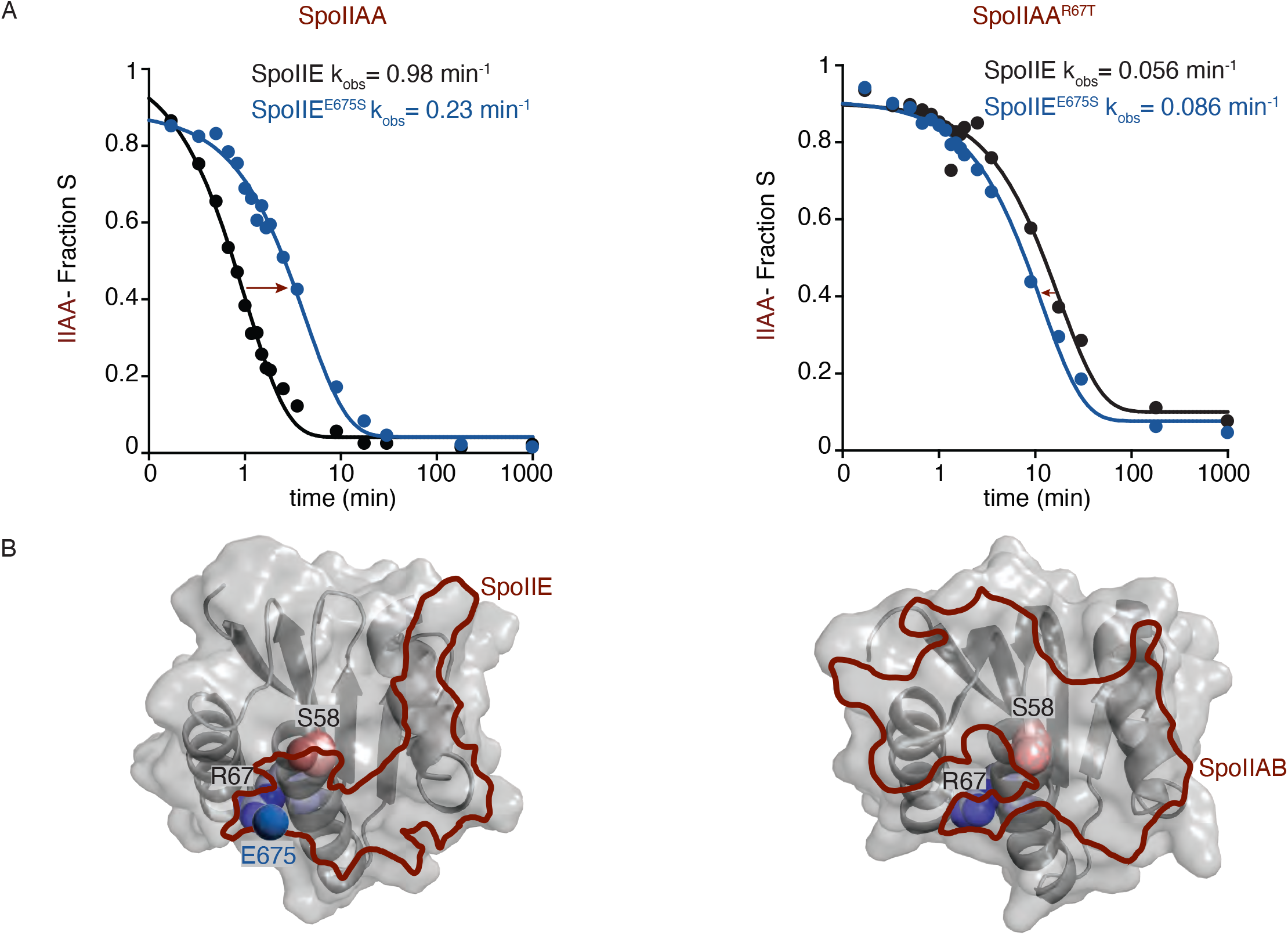
Location of substrate residue complements phosphatase specificity. **A.** Rate of SpoIIAA-P and SpoIIAA^R67T^ dephosphorylation by SpoIIE and SpoIIE^E675S^ over time. Plot on the *left* measures the fraction of SpoIIAA-P dephosphorylated over time by SpoIIE (black) and SpoIIE^E675S^ (blue) and was fit to an exponential decay function. The measured k_obs_ were SpoIIE 0.98± 0.05 min^-1^ and SpoIIE^E675S^ 0.23± 0.015 min^-1^. The *right* plot measures the fraction of SpoIIAA^R67T^ dephosphorylated over time by SpoIIE (gray) and SpoIIE^E675S^ (light blue) and were fit to an exponential decay function. The k_obs_ was SpoIIE 0.056± 0.0061 min^-1^ and SpoIIE^E675S^ 0.086± 0.0062 min^-1^. Reactions were single-turnover reactions using 0.1 µM SpoIIE, 0.5 µM SpoIIAA-P, 10 mM MnCl_2_, and the error is the error of the fit. **B.** AlphaFold2 structure of SpoIIAA showing the SpoIIE and SpoIIAB binding interface outline. The SpoIIAA structure on the *left* shows the phosphorylation site, S58 (red and light pink), and the residue R67 (blue, and light blue). The SpoIIE binding interface is outlined (red) based on a 1.4 Å probe radius, overlapping both S58 and R67. SpoIIE residue E675 (blue) is in proximity to R67. On the *right*, the SpoIIAB/SpoIIAA binding interface is outlined (red) based on a 1.4 Å probe radius. Phosphorylation site S58 (pink) and R67 (blue and light blue) are notated on the structure.

### Opposing kinases and phosphatases recognize distinct substrate features

One important aspect of phosphatase substrates is that they must first be recognized and phosphorylated by a kinase. We therefore, compared the docking interface of SpoIIAA with SpoIIE and with its cognate kinase, SpoIIAB (for which a co-crystal structure has been determined) (Masuda et al. 2004). The interfaces were substantially non-overlapping, with much more extensive contacts formed between SpoIIAB and SpoIIAA than SpoIIE and SpoIIAA (Fig. 5B). This divergence of substrate interaction interface between kinase and phosphatase suggests that substrate specificity has the potential to evolve independently for each enzyme. However, R67 was part of both interfaces, suggesting some overlap in key specificity determinants.

## Discussion

To control critical transcriptional programs, SpoIIE and RsbU must discriminate between their paralogous substrate proteins with high fidelity. How this specificity is achieved was a mystery because both phosphatases present their active sites on the solvent-exposed surface and have similar K_M_ for both substrates(Ho & Bradshaw 2021). Here we discovered that SpoIIE and RsbU have three variable loops that recognize the cognate substrate protein and facilitate dephosphorylation through a conserved allosteric element. Because related phosphatase/substrate pairs control diverse transcriptional programs across bacterial species, we postulate that this mechanism is generalizable, providing a molecular framework for understanding phosphatase specificity. Below we discuss the implications of this mechanism for the evolution of new signaling pathways and for the broader PPM family of phosphatases.

### Substrate docking interactions control phosphatase activity

We discovered that there are three variable loops on the PPM phosphatase domain that discriminate between substrates through direct interactions: the switch-loop (between α2 and β4), the flap (between β7 and β8), and the α3/4 loop. Substitutions of single amino acids in each loop were sufficient to change substrate preference by as much as tenfold, and combining three substitutions was sufficient for a five-hundred-fold change in specificity. Intriguingly, the effect of these substitutions on specificity was entirely through decreasing the k_cat_ for the cognate substrate and increasing the k_cat_ for the non-cognate substrate. We therefore infer that these docking interactions are critical for positioning the substrate protein and organizing the active site. Future high-resolution experimental structures will be required to reveal the specific structural changes that are required for phosphatase activation. However, we discovered a second class of substitutions that made SpoIIE and RsbU more promiscuous and are buried in the core of the phosphatase domain, interacting with a conserved switch element that controls metal cofactor binding and catalysis. Importantly, the flap and switch-loop directly contact the switch element, suggesting a mechanism for how substrate docking is transduced to control the active site.

### Conservation of substrate docking interactions

Our analysis of PPM phosphatase domain sequences confirmed that the switch-loop, flap, and the α3/4 loop are variable regions across PPM family phosphatases, supporting the possibility that they may be conserved elements for achieving substrate specificity. There is one structure in the PDB of a phosphatase bound to its substrate protein, the *A. thaliana* drought-tolerance response phosphatase, HAB1, bound to its substrate, the kinase SnRK2.6 (Soon et al. 2012) (Fig S8). This structure revealed that W385 of the HAB1 flap is a critical latch for interaction with both SnRK2.6 and the competing abscisic acid receptor PYR1. Similar to SpoIIE^I723^ and RsbU^S251^, they additionally identified flap residues V393 and Y404 that are more proximal to the switch element of HAB1 as making important contacts to SnRK2.6. Our analysis of the structure additionally reveals that HAB1^E323,T324^ in the switch-loop (at the equivalent position of SpoIIE^E675^ and RsbU^S251^) and HAB1^S490^ in the α3/4 loop (at the equivalent position of SpoIIE^P752^ and RsbU^R280^) make contacts to SnRK2.6 but not PYR1. This suggests that these critical features for substrate recognition are conserved across kingdoms despite the evolutionary divergence between the phosphatases and despite the fact that they act on unrelated substrate proteins. Supporting this conclusion, the flap region has also been shown to contribute to the specificity of human PPM phosphatases (Gilmartin et al. 2014; Waschbüsch et al. 2021; Yamaguchi et al. 2005).

### The evolutionary diversification of phosphatases

The canonical evolutionary model for how proteins evolve new substrate specificity is that following a gene duplication event, mutations create a promiscuous intermediate before subsequent mutations block recognition of the original substrate and optimize recognition of the new substrate(Khersonsky & Tawfik 2010). Our findings suggest that PPM phosphatases have a built-in path for this evolution, with mutations in the switch element causing substrate promiscuity and variable loop regions directly interacting with substrates providing discrimination. We speculate that this ordered pathway may underly the diversification of PPM phosphatases, particularly in species of bacteria and plants that have large numbers of PPM phosphatases (often more than fifty) (Zhang & L. Shi 2004; Moorhead et al. 2009).

The evolution of phosphatase specificity is additionally constrained by the fact that each substrate protein is necessarily also the substrate of an opposing protein kinase. For SpoIIAA, the phosphatase/substrate and kinase/substrate interfaces are largely non-overlapping, suggesting that there is significant room for phosphatases and kinases to independently evolve specificity for the same substrate proteins. Whether these same principles hold for other kinase/phosphatase pairs will be of significant interest as the mechanism of substrate discrimination is uncovered for more phosphatases.

## Materials and Methods

### Strain construction

Strains were grown in liquid Lennox lysogeny broth (LB, Sigma Aldrich) or on plates supplemented with 15% Bacto agar (Difco). Competence medium (MC) was used during Bacillus transformation. Antibiotics were added where appropriate to select for plasmids and for transformant selection but strains with genomic markers were not generally grown on selective media. Isopropyl-beta-D-thiogalactoside (IPTG) was used at 1 mM, 5-bromo-4-chloro-3-indolyl-beta-D-galacto-pyranoside (X-gal) was used at 80 µg/ml. Standard molecular biology techniques were used to construct DNA plasmids using isothermal assembly (Gibson cloning) to generate new constructs and Quikchange mutagenesis for site directed mutagenesis. Table 1 provides all strains and Table 2 provides all primers used in this study.

All strains were constructed in the *Bacillus subtilis* PY79 strain background. To make the σ^F^ reporter strain, genomic DNA from *B. subtilis* strains containing *ΔrsbR rsbS rsbT rsbU rsbV rsbW sigB rsbX::kan* (Carniol, Kim, et al. 2004), *amyE::pspoIIQ-lacZ cm* (Boylan et al. 1992) and *spoIIAB::spoIIAB^R105C^ spec* (Carniol, Eichenberger, et al. 2004) genetic modifications and were sequentially transformed into the *ΔspoIIE::phleo* (Carniol, Eichenberger, et al. 2004) parent strain. Transformants were selected based on antibiotic resistance and the ability to break down starch. The *spoIIAB^R105C^ spec* construct was introduced by long flanking homology PCR and its sequence was confirmed by colony PCR with primers flanking the spoIIAA operon for sequencing. The σ^F^ expressing strain for testing viability during vegetative growth was generated by sequentially transforming a *B. subtills* strain with *ywrk::Tn917::amyE::pspank-spoIIAoperon tet* (Bradshaw & Losick 2015) with genomic DNA containing *ΔrsbR rsbS rsbT rsbU rsbV rsbW sigB rsbX::kan*, *amyE::pSpoIIQ-lacZ cm*, and *ΔspoIIE::phleo*. The σ^B^ reporter strains were constructed as described previously (Ho & Bradshaw 2021).

### σ^F^ Toxicity assay

Strains were grown in LB/MLS to an OD600 of approximately 0.35. Cells were ten-fold serially diluted and spotted on LB, MLS, and x-gal, with and without IPTG at 37°C overnight. Another aliquot of the culture was plated on LB/MLS, and a Whatman paper disc approximately 1 cm that was saturated with 50 µL of 1M IPTG was added to the center of the plate after spreading the cells. The cells were grown at 37°C overnight.

### Genetic screen

*B. subtilis* genomic DNA containing RsbT/RsbU was amplified by PCR using GoTaq DNA polymerase mix for thirty cycles without modification from the manufacturers protocol (Promega). DNA sequencing revealed that inserts had on average one to two mutations per product. Pools of the mutagenized PCR product were assembled into the pHB201 digested vector using isothermal (Gibson) assembly and transformed into E. coli DH5α cells. *E. coli* cells were pooled, plasmid DNA was extracted and transformed into the *Bacillus* screen strain using natural competence. *B. subtilis* cells were plated on Difco sporulation medium (DSM) containing MLS, IPTG, and X-gal. Blue colonies were selected and restruck on DSM plates with and without IPTG, as well as LB/MLS plates. Plasmids from colonies that retested as σ^F^ positive were isolated and inserts were sequenced by Sanger sequencing.

### Protein expression and purification

All proteins were expressed in *E. coli BL21 (DE3)* cells grown at 37°C to an OD600 of 0.4 and induced at 16°C for 14-18 hours with 1 mM (IPTG). Cells were harvested and purified as follows (protocols adapted with modification from (Baral et al. 2024; Ho & Bradshaw 2021; Bradshaw et al. 2017)): *SpoIIE and SpoIIE variants*: Cell pellets were resuspended in lysis buffer with 1mM phenylmethylsulfonyl fluoride (PMSF) (50 mM K•HEPES, pH 8, 200 mM NaCl, 20 mM imidazole, 10% glycerol, 0.5 mM dithiothreitol (DTT)) and were lysed using three passes in a microfluidizer at 10,000 PSI. Cell lysates were cleared by spinning at 16,000 RPM for 30-45 min in an Avanti JA-20 rotor. Cleared lysates were then bound to Ni-NTA resin (2ml/L of culture) on the column by gravity flow after the Ni-NTA resin slurry was equilibrated with lysis buffer. The resin was then washed with 10 CV of lysis buffer containing 20 mM imidazole and eluted with 200 mM imidazole. The 6-His tag was cleaved with 3C protease in dialysis with lysis buffer at 4°C overnight. The 6-His cleaved tag and 3C protease was subtracted by passing over the Ni-NTA resin. The protein was spin-concentrated before the gel filtration run. It was further purified on a Superdex 75 16/60 column that was equilibrated in 20mM K•HEPES, pH 8, 50 mM NaCl, 2 mM DTT on the AKTA FPLC. The fractions were pooled, concentrated approximately between 46-200 µM, flash-frozen and stored at -80°C.

*SpoIIAA and SpoIIAA variants*: Cell pellets were resuspended in lysis buffer with 1mM phenylmethylsulfonyl fluoride (PMSF) (50 mM K•HEPES, pH 8, 100 mM NaCl, 20 mM imidazole, 10% glycerol, 0.5 mM DTT) and were lysed using two passes in a microfluidizer at 10,000 PSI. Cell lysates were cleared by spinning at 16,000 RPM for 30 min in a Sorvall SS-34 rotor. Cleared lysates were run over a HisTrap HP column on an AKTA FPLC. Fractions were pooled and the 6-His tag was cleaved with 3C protease in dialysis with lysis buffer at 4°C overnight. The 6-His cleaved tag and 3C protease were subtracted by passing over equilibrated Ni-NTA resin. Protein was spin-concentrated before the gel filtration run. It was further purified on a Superdex 75 16/60 column that was equilibrated in 50 mM K•HEPES, pH 8, 100 mM NaCl, 10% glycerol and 2 mM DTT on the AKTA FPLC. The fractions were pooled, concentrated approximately between 270-500 µM, flash-frozen and stored at -80°C.

*SpoIIAA-P*: Cell pellet was resuspended in lysis buffer with 1 mM PMSF (50 mM K•HEPES, pH8, 100 mM NaCl, 20 mM imidazole, 10% glycerol, 0.5 mM DTT) and was lysed using two passes on a microfluidizer at 10,000 PSI. Cell lysate was cleared by spinning at 16,000 RPM for 30 min in an Avanti JA-20 rotor. Cleared lysate was then bound to Ni-NTA resin (2ml/L of culture) on the column by gravity flow after the Ni-NTA resin slurry was equilibrated with lysis buffer. The resin was then washed with 10 CV of lysis buffer containing 20 mM imidazole and eluted with 200 mM imidazole. The 6-His tag was cleaved with 3C protease in dialysis with lysis buffer at 4°C overnight. The 6-His cleaved tag and 3C protease was subtracted by passing over the Ni-NTA resin. The protein was spin-concentrated before the gel filtration run. It was further purified on a Superdex 200 column that was equilibrated in 50 mM K•HEPES, pH 8, 100 mM NaCl, 10% glycerol, 2 mM DTT. The fractions were pooled, concentrated approximately 200-300 µM, flash-frozen, and stored at -80°C.

*RsbV*: BL21 (DE3) cells were transformed with pET47bRsbV each time before the protein was expressed. Cell pellets were resuspended in lysis buffer with 200uM PMSF (50 mM K•HEPES, pH8, 200 mM NaCl, 20 mM imidazole, 10% glycerol, 0.5 mM DTT), and were lysed using two passes on a microfluidizer at 10,000 PSI. Cell lysate was cleared by spinning at 16,000 RPM for 45 min in an Avanti JA-20 rotor. Cleared lysates were then bound to Ni-NTA resin (2ml/L of culture) on the column by gravity flow after the Ni-NTA resin slurry was equilibrated with lysis buffer. The resin was then washed with 10 CV of lysis buffer containing 20 mM imidazole and eluted with 200 mM imidazole. The 6-His tag was cleaved with 3C protease in dialysis with lysis buffer at 4°C overnight. The 6-His cleaved tag and 3C protease was subtracted by passing over the Ni-NTA resin. The protein was spin-concentrated before the gel filtration run. It was further purified on a Superdex 75 16/60 column that was equilibrated in 50 mM K•HEPES, pH 8, 100 mM NaCl, 10% glycerol, 2 mM DTT. The fractions were pooled, concentrated approximately 150 µM, flash-frozen and stored at -80°C.

*SpoIIAB:* Cell pellets were resuspended in lysis buffer with 1mM phenylmethylsulfonyl fluoride (PMSF) (50 mM K•HEPES, pH 7.5, 200 mM NaCl, 10 mM MgCl_2_, 20 mM imidazole, 10% glycerol, 0.5 mM DTT) and were lysed using two passes in a microfluidizer at 10,000 PSI. Cell lysates were cleared by spinning at 16,000 RPM for 30 min in a Sorvall SS-34 rotor. Cleared lysates were run over a HisTrap HP column on an AKTA FPLC. Fractions were pooled and the protein was spin-concentrated before the gel filtration run. It was further purified on a Superdex 75 16/60 column that was equilibrated in 50 mM K•HEPES, pH 7.5, 175 mM NaCl, 10% glycerol, 10 mM MgCl2 and 1 mM DTT on the AKTA FPLC. The 6H-sumo tag was left uncleaved to aid with removal during phosphorylation reactions. The fractions were pooled, concentrated to approximately 200 µM, flash-frozen and stored at -80°C.

### Phosphatase assays

Phosphatase assays were performed with SpoIIAA that was labeled with ^32^P by incubating SpoIIAA (45 µM), 6His-sumo-SpoIIAB (55 µM), 50 µCi of γ-^32^P ATP overnight at room temperature in 50 mM K•HEPES, pH 7.5, 50 mM KCl, 2 mM DTT, 0.75 mM MgCl_2_. Unincorporated nucleotide was removed by buffer exchange using a Zeba spin column (Pierce) equilibrated with 25 mM K•HEPES, pH 7.5, 200 mM NaCl. 6H-sumo-SpoIIAB was removed by incubating it with Q-Sepharose resin equilibrated in 50 mM K•HEPES, pH 7.5, 50 mM KCl, 2 mM DTT, 0.75 mM MgCl_2_. The flowthrough fraction from the Q Sepharose resin containing SpoIIAA-^32^P was then exchanged into 50 mM K•HEPES, pH 8, 100 mM NaCl using Zeba spin column to remove an unincorporated nucleotide and free phosphate. The labeled SpoIIAA-^32^P was aliquoted and frozen at -80°C for future use.

To produce ^32^P-labeled RsbV-P, 40 µM RsbV, 45 µM 6His-RsbW and 100 µCi of γ-^32^P ATP were incubated overnight at room temperature in 50 mM K•HEPES, pH7.5, 50 mM KCl, 10 mM MgCl2, and 2 mM DTT. Unincorporated nucleotide was removed by buffer exchange using Zeba column equilibrated in 50 mM K•HEPES, pH 8, 100 mM NaCl. 6His-RsbW was then removed by Ni-NTA resin equilibrated in 50 mM K•HEPES, pH 8, 100 mM NaCl, 20 mM imidazole. The RsbV-^32^P flow-through fraction from the Ni-NTA resin was then buffer exchanged into 50 mM K•HEPES, pH 8, 100 mM NaCl buffer using three sequential Zeba spin columns to remove all unincorporated nucleotide and free phosphate. The labeled RsbV-^32^P was aliquoted and frozen at -80°C.

All phosphatase assays were conducted at room temperature in 50 mM K•HEPES, pH8, 100 mM NaCl. The concentrations of enzyme, substrate and metal cofactor (MnCl_2_ or MgCl_2_) were varied as indicated. SpoIIAA reactions had 0.2 mg BSA added to prevent protein adhesion to reaction tubes. Reactions were stopped with 0.5 M EDTA, pH 8, and 2% SDS, then run on PEI-Cellulose TLC plates that were developed in 1 M LiCl2 and 0.8 M acetic acid. The plates were imaged on an Amersham typhoon scanner and quantified with ImageQuant. The error reported is from the error of the fit.

### p-Nitrophenyl Phosphate (PNPP)

PNPP assay was conducted at room temperature by mixing 50 mM HEPES, pH 8, 100 mM NaCl, 0.5 uM of enzyme, 20 mM MnCl2, and increasing concentrations of PNPP (0.5 mM-25 mM) in a 96 well plate. Reactions were started with PNPP and hydrolysis of PNPP to p-nitrophenol was measured at 405 nm in a plate reader.

### AlphaFold2 structure predictions

AlphaFold2 predictions were performed using ColabFold (Mirdita et al. 2022).

Alphafold2_multimer_v2 was used in unpaired_paired mode with no templates with 3 recycles, 200 iterations, and greedy pairing strategy. The predicted aligned error plots for all AlphaFold2 structures are shown in Fig. S7.

Supplementary Table S1:

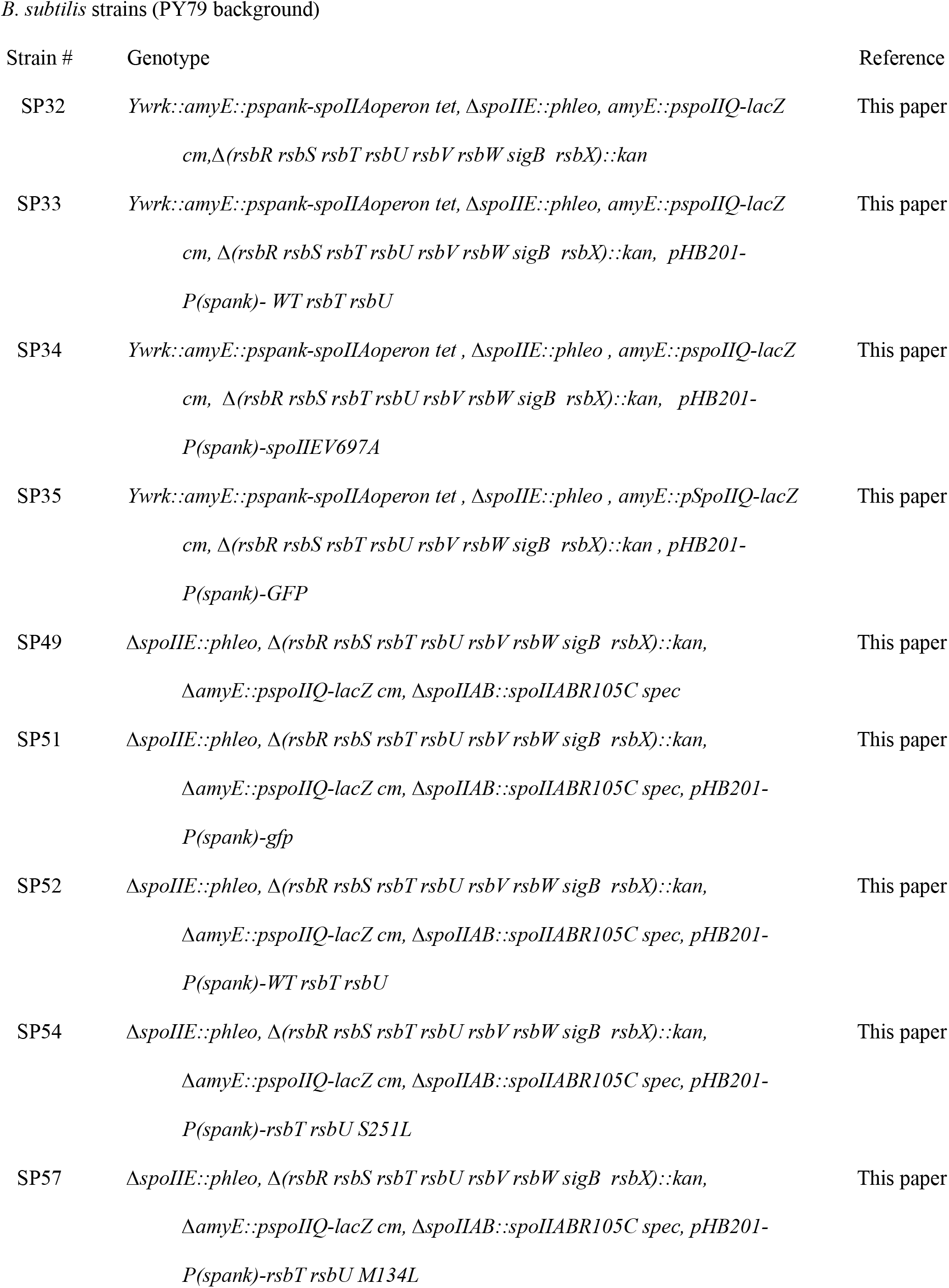
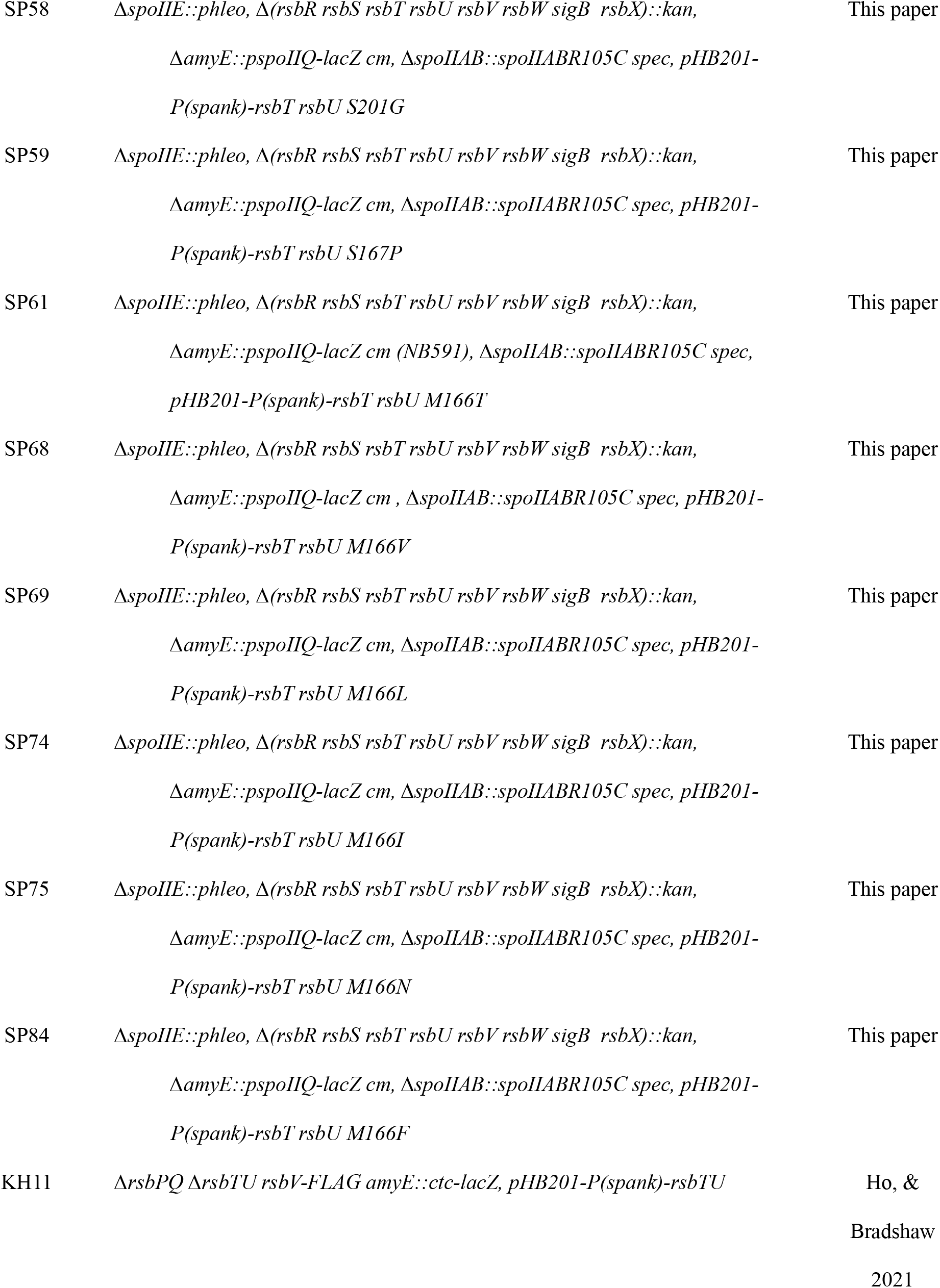
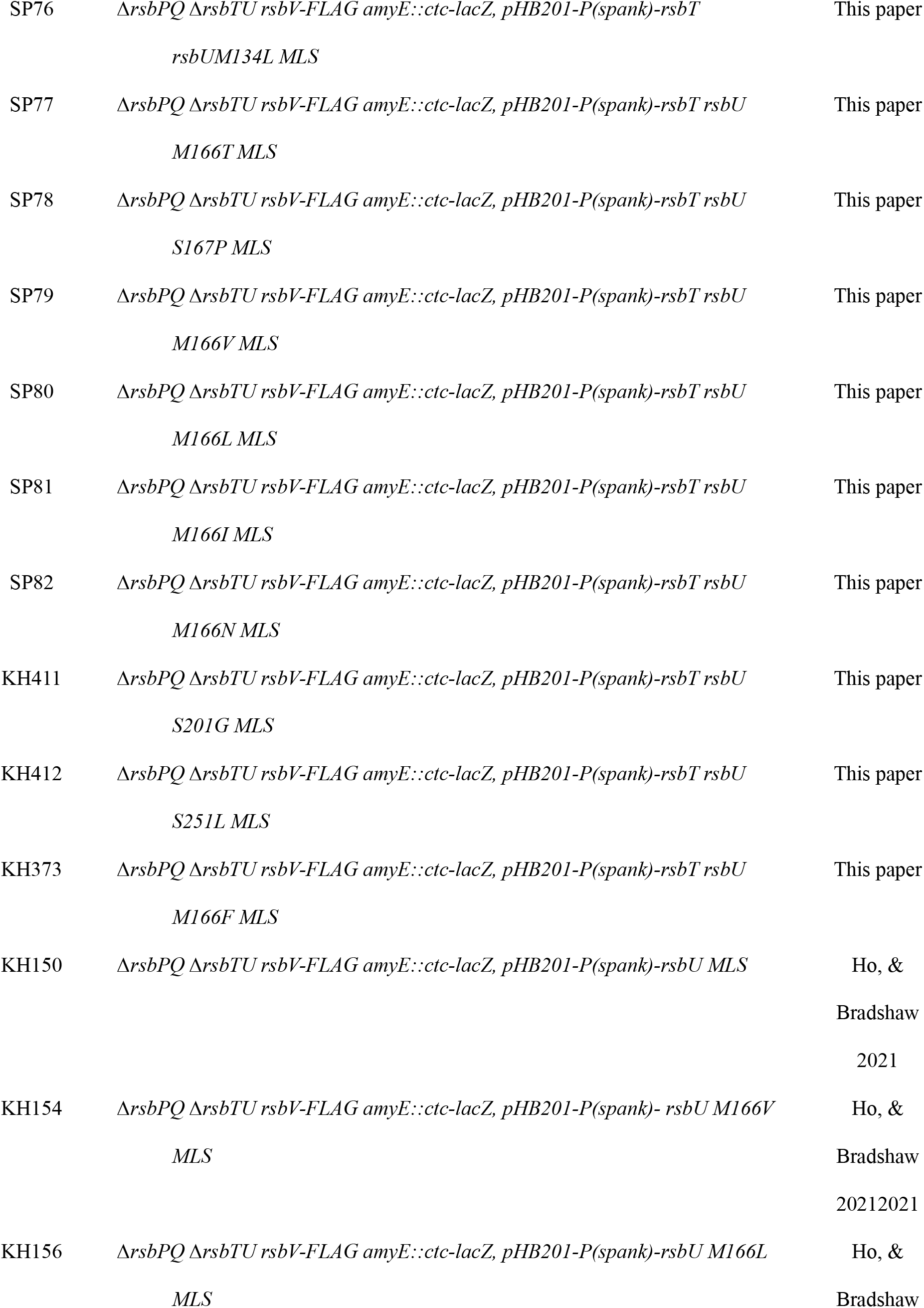
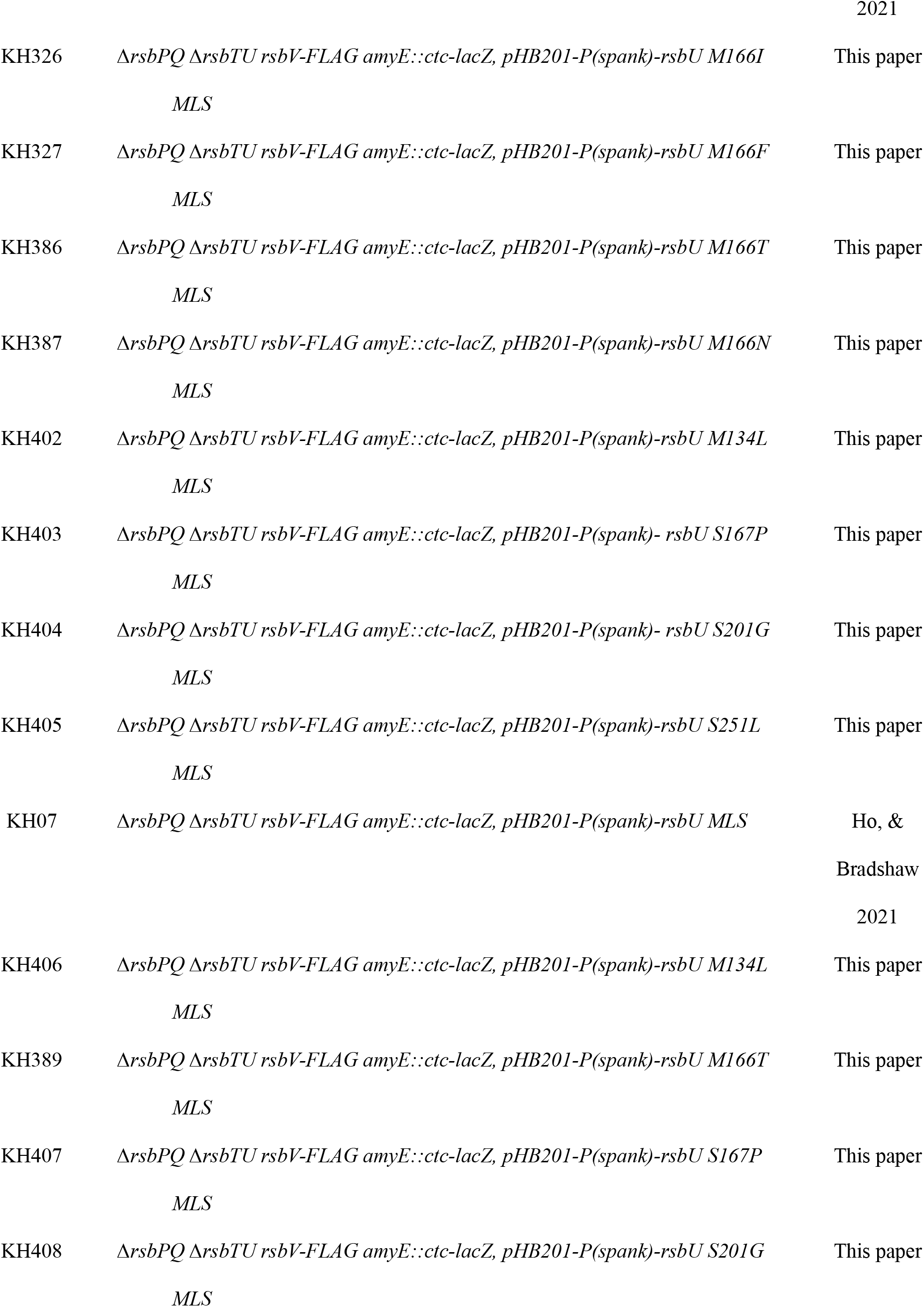
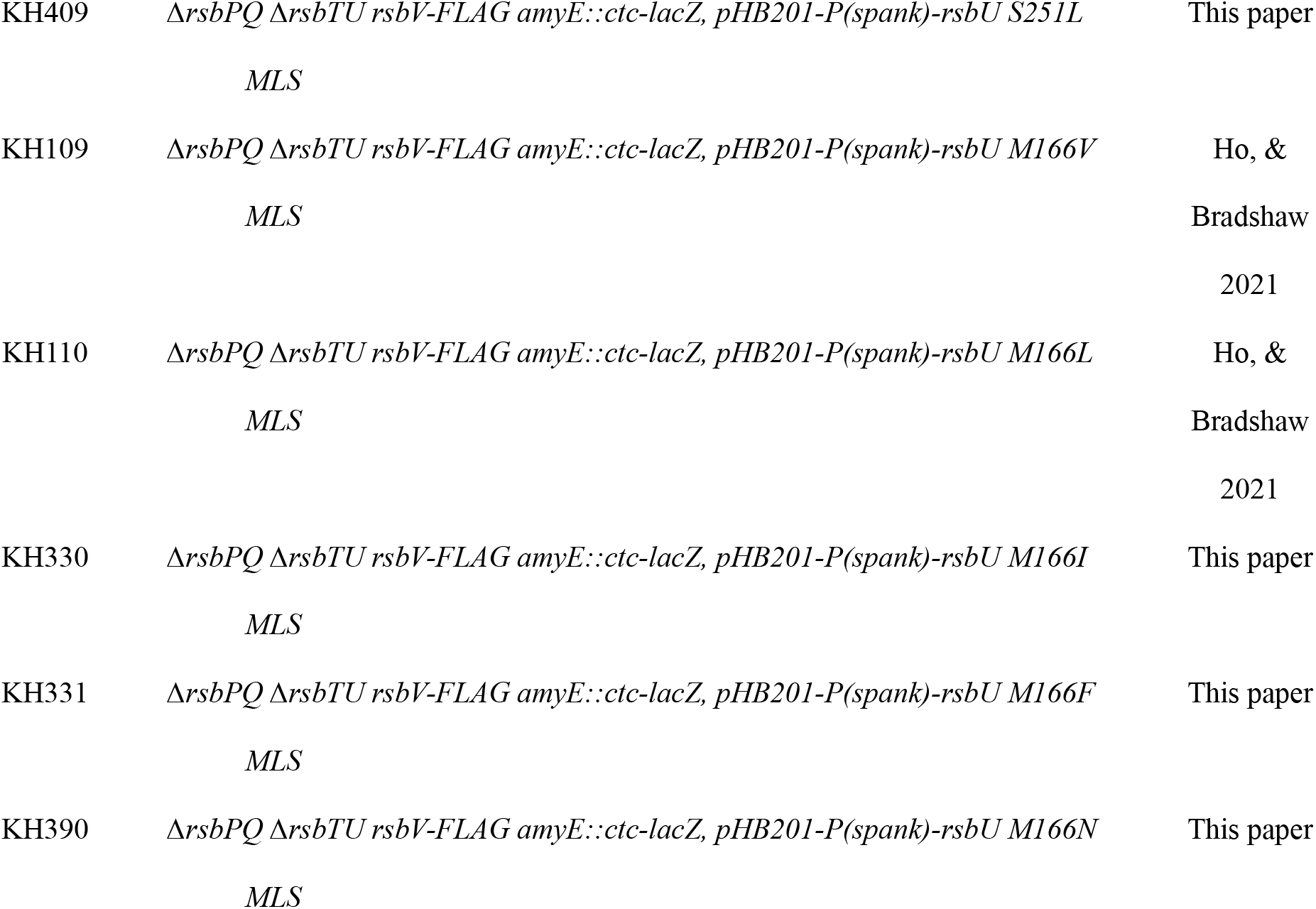

*Supplemental table 2:*

*Supplemental table 2:* *E. coli* strains (BL21 background)
| Strain number | Genotype | Reference |
| --- | --- | --- |
| SP85 | *BL21 (DE3) pET47b-6H-3C-spoIIE-590-827 P752R Kan* | This paper |
| SP86 | *BL21 (DE3) pET47b -6H-3C-spoIIE-590-827 E675S Kan* | This paper |
| SP87 | *BL21 (DE3) pET47b -6H-3C-spoIIE-590-827 I723S Kan* | This paper |
| SP90 | *BL21 (DE3) pET47b 6H-3C-spoIIE-590-827 P752R-E675S-I723S Kan* | This paper |
| NB1732 | *BL21 (DE3) pET47b 6H-3C-spoIIE 590-827 Kan* | Ho, &<br>Bradshaw<br>2021 |
| NB1959 | *BL21 (DE3) pET47b 6H-3C-spoIIAA Kan* | Ho, &<br>Bradshaw<br>2021 |
| SP12 | *BL21 (DE3) pET47b 6H-3C-spoIIAA R67T Kan* | This paper |
| NB781 | *BL21 (DE3) pET47b 6H-3C-spoIIAA spoIIAB Kan* |  |
| NB182 | *BL21 (DE3) pET23a 6H-Sumo-spoIIAB amp* | Ho, &<br>Bradshaw<br>2021 |
| NB1835 | *BL21 (DE3) pET47b 6H-3C-rsbV Kan* | Ho, &<br>Bradshaw<br>2021 |

**Supplemental Figure 1:**
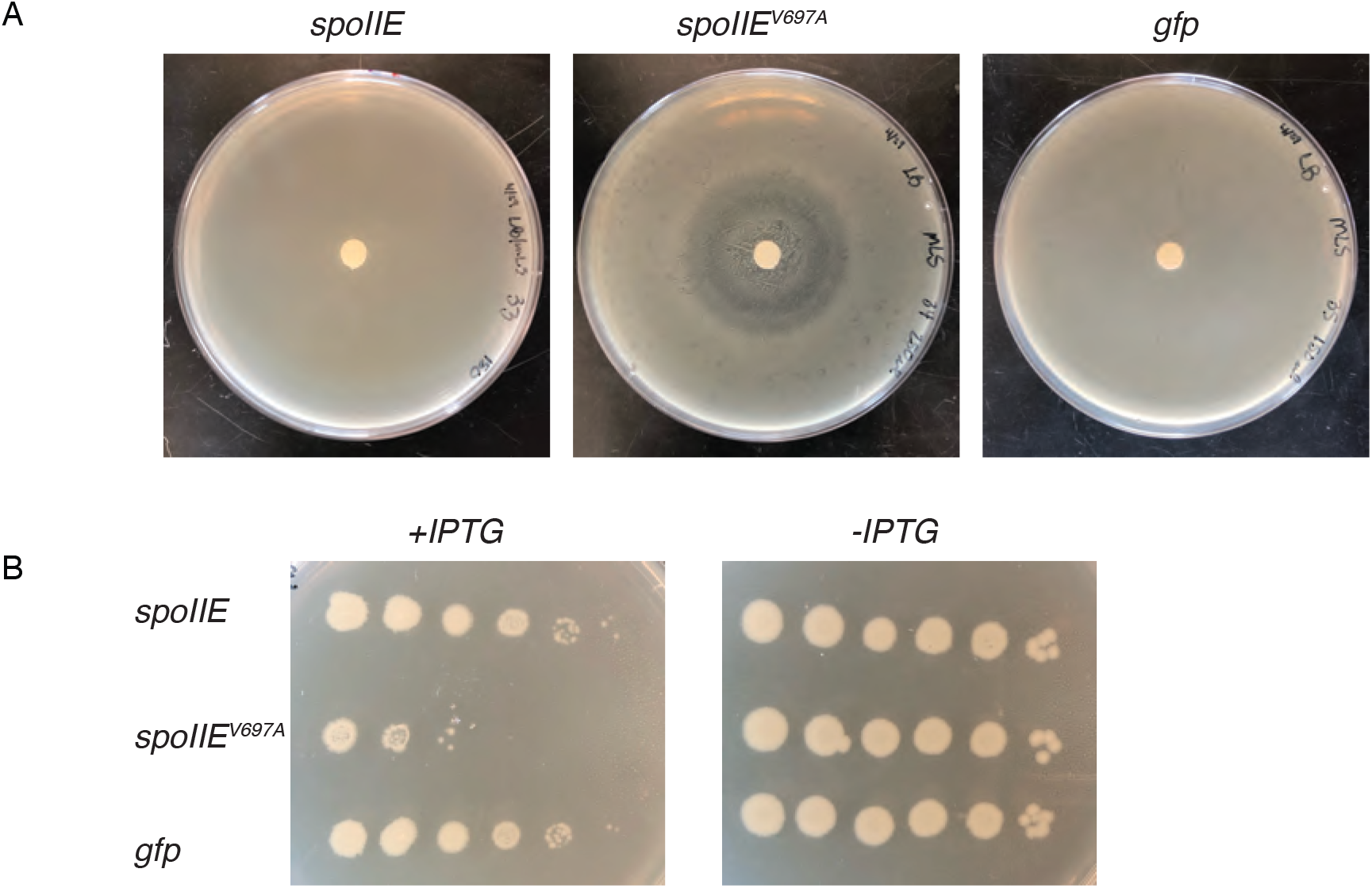
Activation of σ^F^ during vegetative growth is toxic to cells. **A.** Strains with σ^F^ expression under nonsporulating conditions with a *lacZ* reporter for σ^F^ activity. Strain expressing either *spoIIE* (left plate) *spoIIE^V697A^* (middle plate) or *gfp* (right plate) from plasmid pHB201 were grown on LB/MLS plates. A Whatman paper disk of approximately 1cm was saturated with 1M IPTG and placed in the center of the plate. Plates were incubated at 37°C overnight. **B.** Strain expressing either *spoIIE* (top) *spoIIE^V697A^* (middle) or *gfp* (bottom) from plasmid pHB201 were serial diluted 10-fold and grown on LB/MLS plates, plus (left) and minus (right) 1 mM IPTG.

**Supplemental Figure 2:**
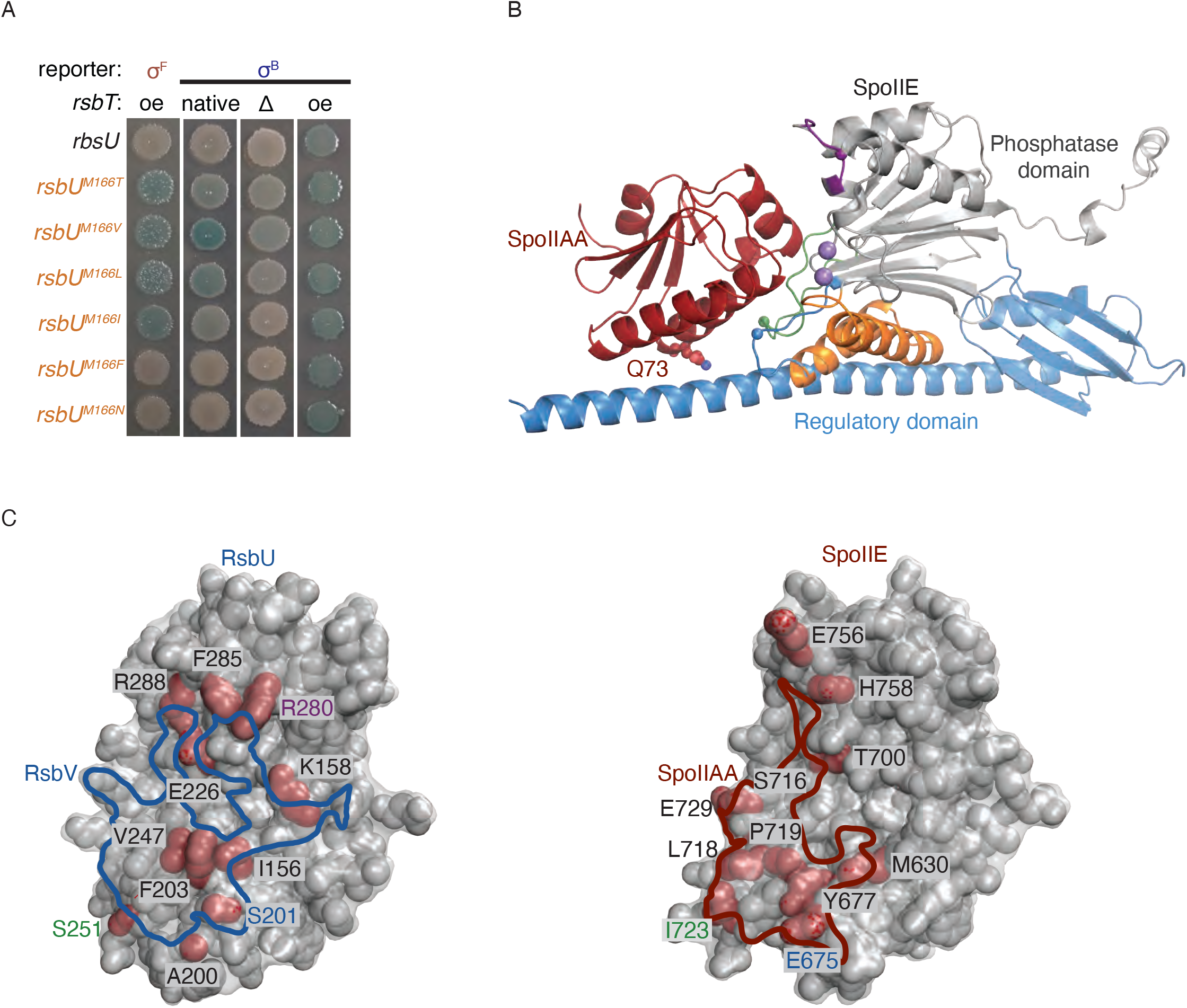
Conservation of phosphatase/substrate recognition. **A.** Strains carrying plasmid expressing M166X mutation located in the switch of RsbU. Reporter strains have *lacZ* under the control of either σ^F^ (left row) or σ^F^ (last three rows) promoter. *rsbT* is overexpressed in the σ^F^ reporter strain. *rsbT* in the σ^B^ reporter strains is either expressed from the native locus (second column from left), deleted (third column from left), or overexpressed (last column). M166X mutation series are M166T, M166V, M166L, M166I, M166F, and M166N. **B.** AlphaFold2 Structure of *B. subtilis* SpoIIE monomer and SpoIIAA showing contact interface between the two proteins. The phosphatase domain of SpoIIE (gray), along with the elements located in the phosphatase domain, switch (orange), flap (green), 2/4 loop (blue), and the 3/4 loop, are noted on the structure. The metal ions (purple) are visible in the active site. Residues E675S (blue), and I723 (green), are represented by spheres within the α1/ β4 loop and flap, respectively. Position of the SpoIIE regulatory domain (blue) in relation to SpoIIAA (red). SpoIIAA shows the position of residue Q73 (red and blue spheres) in relation to the regulatory domain of SpoIIE. **B.** AlphaFold2 structure of *B. subtilis* RsbU (left) and SpoIIE (right). The phosphatase domain (gray) with non-conserved residues (red) whose side chains contact the side chains of RsbV and SpoIIAA, respectively. The RsbU/RsbV binding interface is outlined in blue, and SpoIIE/SpoIIAA in red based on 1.4 Å probe radius.

**Supplemental Figure 3:**
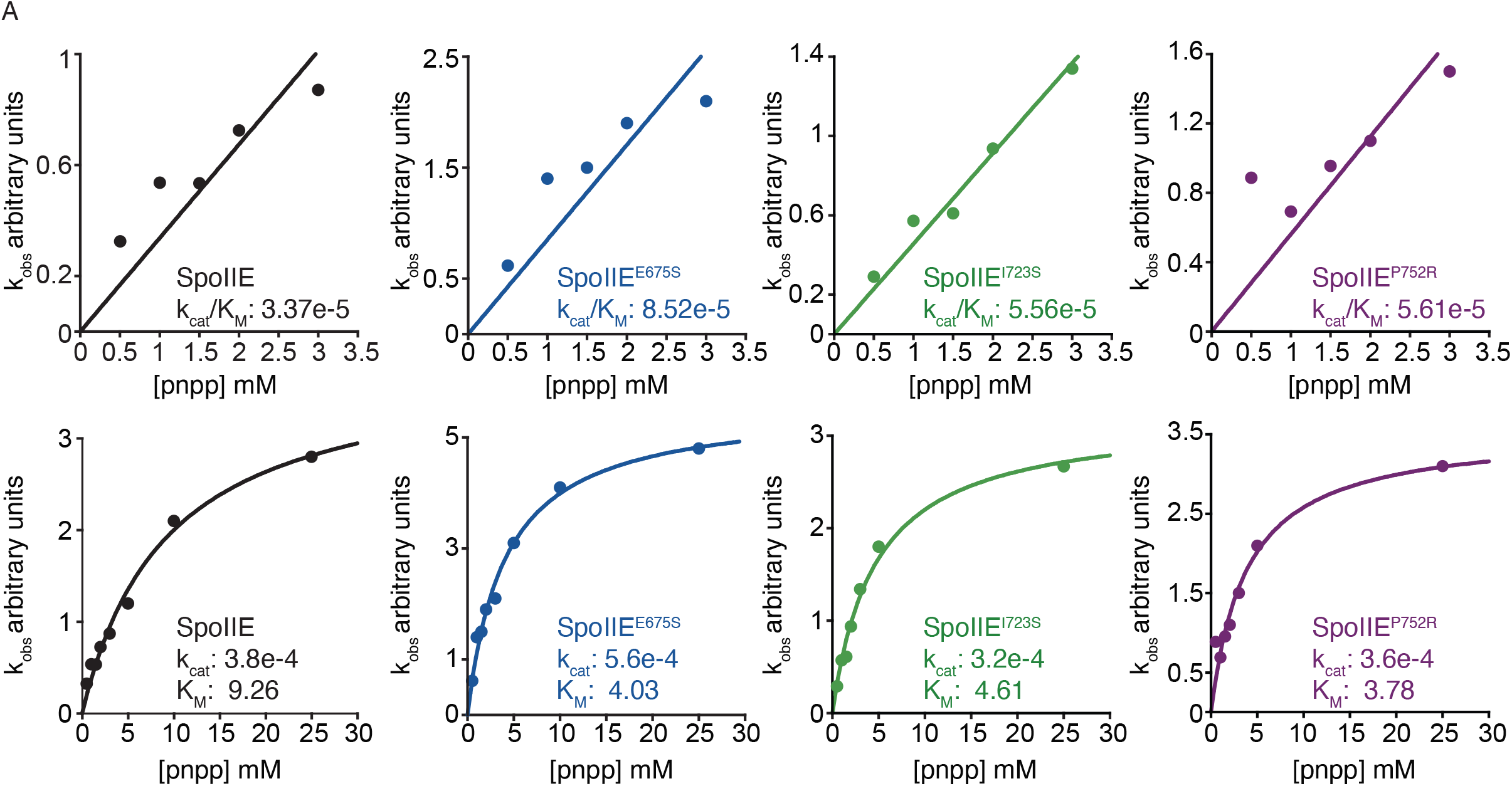
Changes in phosphatase activity are substrate-specific. **A.** The rate of p-nitrophenyl phosphate dephosphorylation during the initial linear phase was measured for SpoIIE, SpoIIE^E675S^, SpoIIE^I723S^, and SpoIIE^P752R^. Observed velocities (in arbitrary absorbance units per time) were plotted as a function of pnpp concentration. Top plots were fit to the linear equation (in KaleidaGraph) (k_cat_/K_M_)*[pnpp]. The lower plots were fit to the Michaelis-Menten equation (in KaleidaGraph) k_cat_*[pnpp]/(K_M_ + [pnpp]).

**Supplemental Figure 4:**
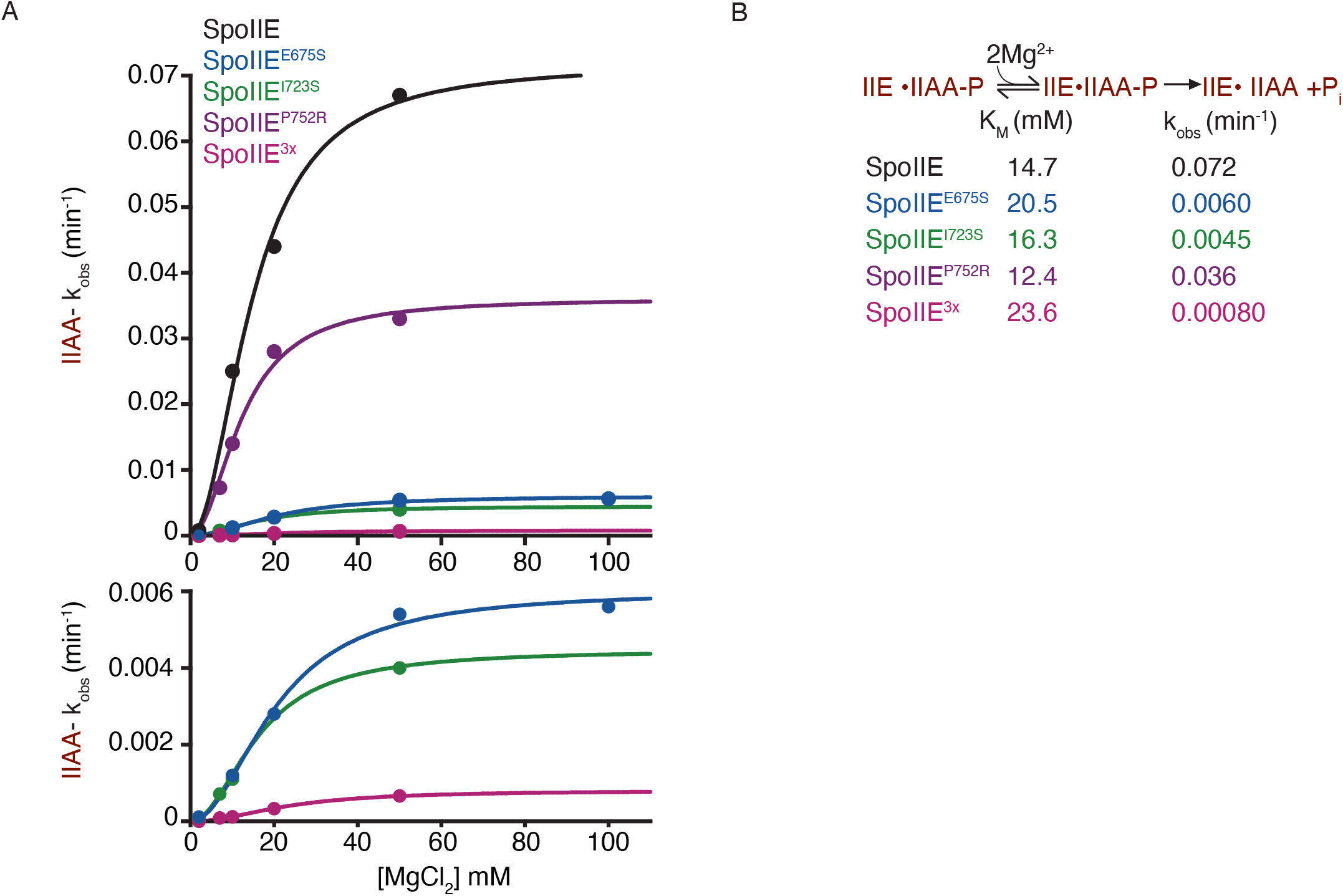
Recruitment of metal cofactor is not impacted. **A.** Activity of SpoIIE (black), SpoIIE^E675S^ (blue), SpoIIE^I723S^ (green), SpoIIE^P752R^ (purple), and SpoIIE^3x^ (pink) as a function of MgCl_2_ concentration with SpoIIAA. The data plotted was fit to a cooperative model using the equation k_obs_=k_cat_*[MgCl_2_]^2^/(K_M_^2^+[MgCl_2_]^2^). The measured K_M_ were SpoIIE 14.7± 1.2 µM, SpoIIE^E675S^ 20.5± 1.4 µM, SpoIIE^I723S^ 16.3± 0.81 µM, SpoIIE^P752R^ 12.4± 1.0 µM and SpoIIE^3x^ 23.6± 1.2 µM. The error is the error of the fit. The reactions were single turnover reactions with varying concentrations of MgCl_2_, 1µM SpoIIE, and 0.05 SpoIIAA. **B.** Graphic scheme showing a summary of the dephosphorylation reaction of SpoIIAA by SpoIIE.

**Supplementary Figure 5:**
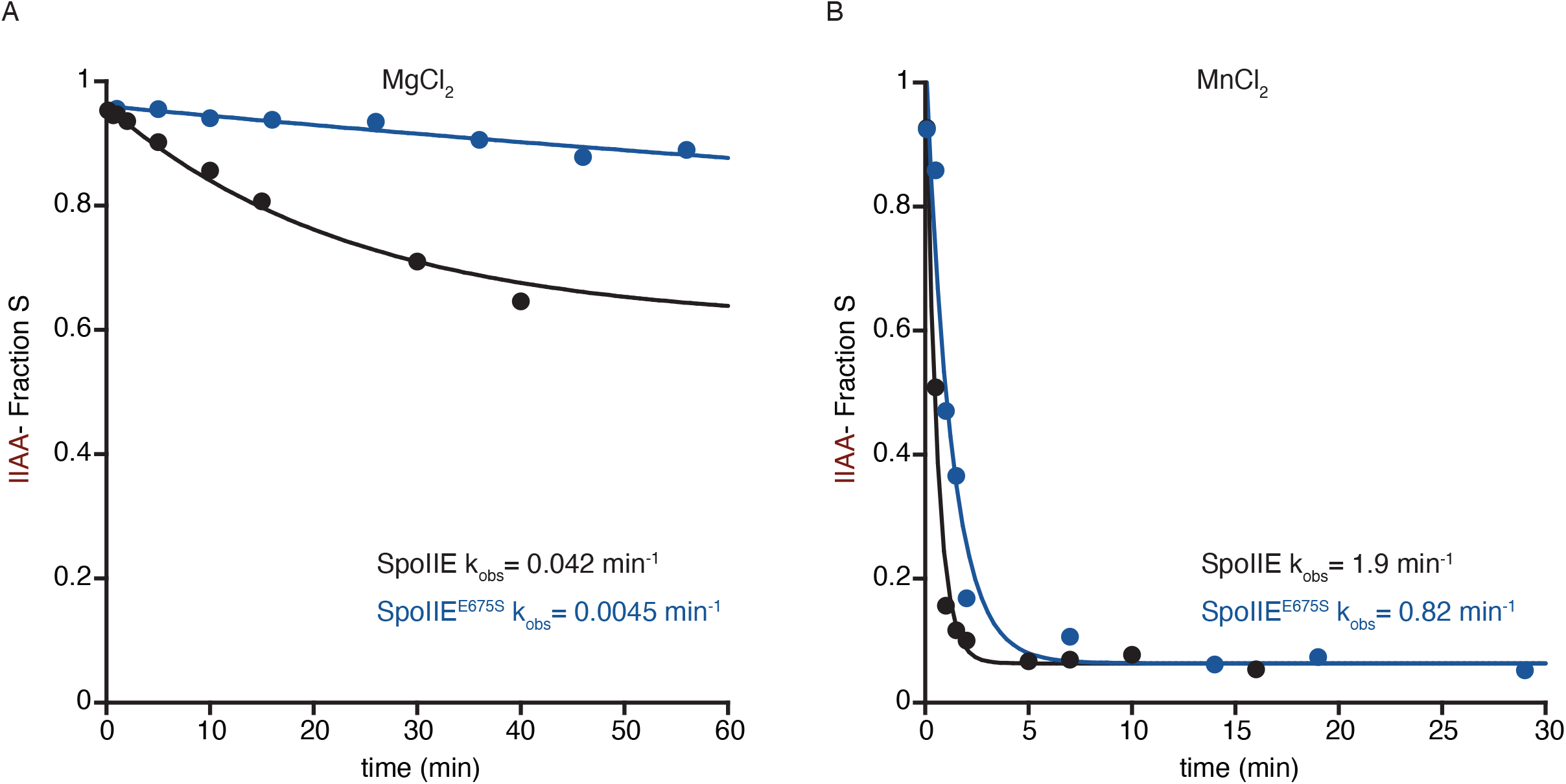
Metal cofactor influences catalysis of cognate substrate. Activity of SpoIIE (black) and SpoIIE^E675S^ (blue) with SpoIIAA with either MgCl_2_ or MnCl_2_. Reactions measured the fraction of SpoIIAA as it decreased over time. **A**. Single turnover reactions were conducted with 1 µM SpoIIE, 0.5 µM SpoIIAA-P, and 10 mM MgCl_2_. The k_obs_ were SpoIIE 0.042± 0.0023 min^-1^ and SpoIIE^E675S^ 0.0045± 0.00036 min^-1^. The data was fit to an exponential decay function. The error is the error of the fit. **B.** Single turnover reactions were conducted with 1 µM SpoIIE, 0.5 µM SpoIIAA-P, and 10 mM MnCl_2_. The k_obs_ were SpoIIE 1.9± 0.18 min^-1^and SpoIIE^E675S^ 0.82± 0.14 min^-1^. The data was fit to an exponential decay function. The error is the error of the fit.

**Supplemental Figure 6:**
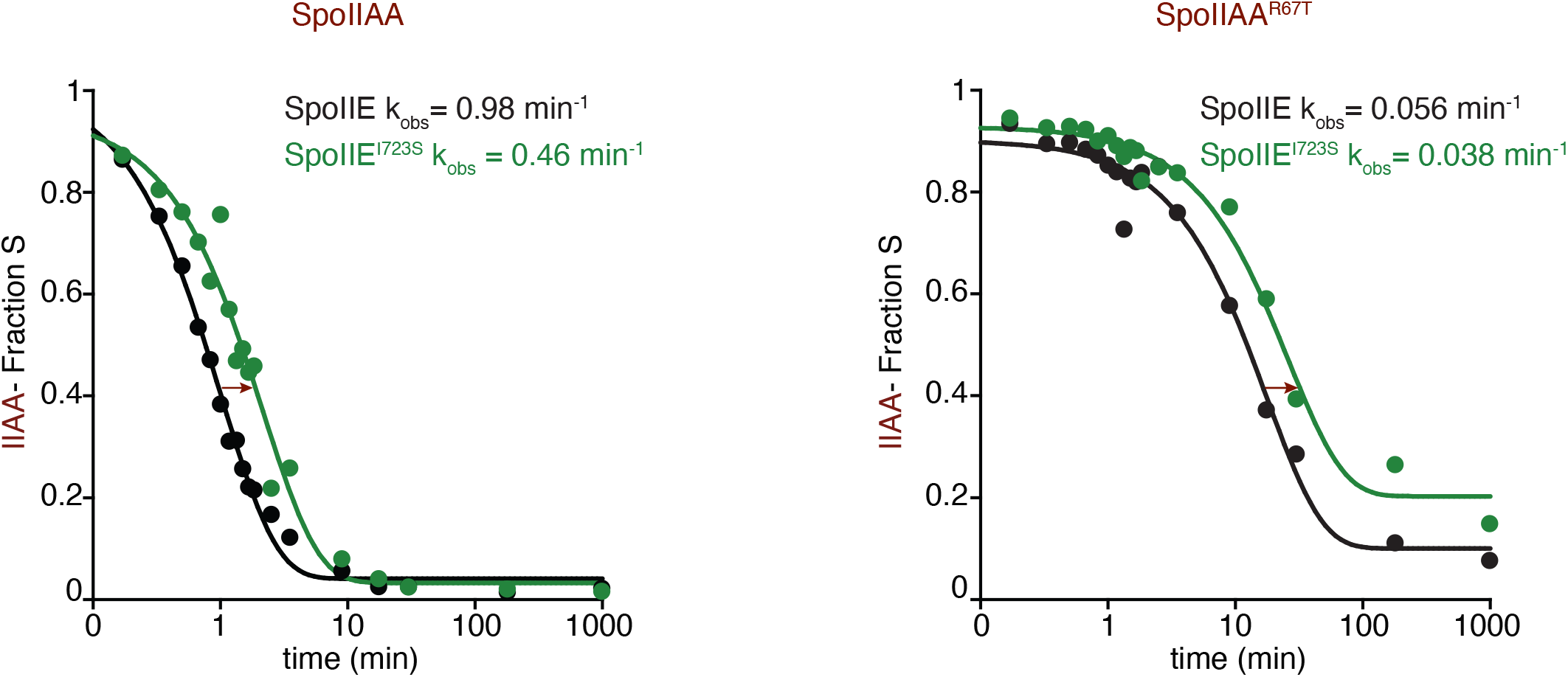
SpoIIE^I723S^ does not complement SpoIIAA^R67T^. **A.** Rate of SpoIIAA-P and SpoIIAA^R67T^ dephosphorylation by SpoIIE and SpoIIE^I723S^ over time. The plot on the *left* measures the fraction of SpoIIAA-P dephosphorylated over time by SpoIIE (black) and SpoIIE^I723S^ (green) and was fit to an exponential decay function. The measured k_obs_ were SpoIIE 0.98± 0.05 min^-1^ and SpoIIE^I723S^ 0.46± 0.049 min^-1^. The *right* plot measures the fraction of SpoIIAA^R67T^ dephosphorylated over time by SpoIIE (gray) and SpoIIE^I723S^ (light green) and was fit to an exponential decay function. The k_obs_ were SpoIIE 0.056± 0.0061 min^-1^ and SpoIIE^I723S^ 0.038± 0.0039 min^-1^. Reactions were single-turnover reactions using 0.1 µM SpoIIE, 0.5 µM SpoIIAA-P, 10 mM MnCl_2_, and the error is the error of the fit.

**Supplemental Figure 7:**
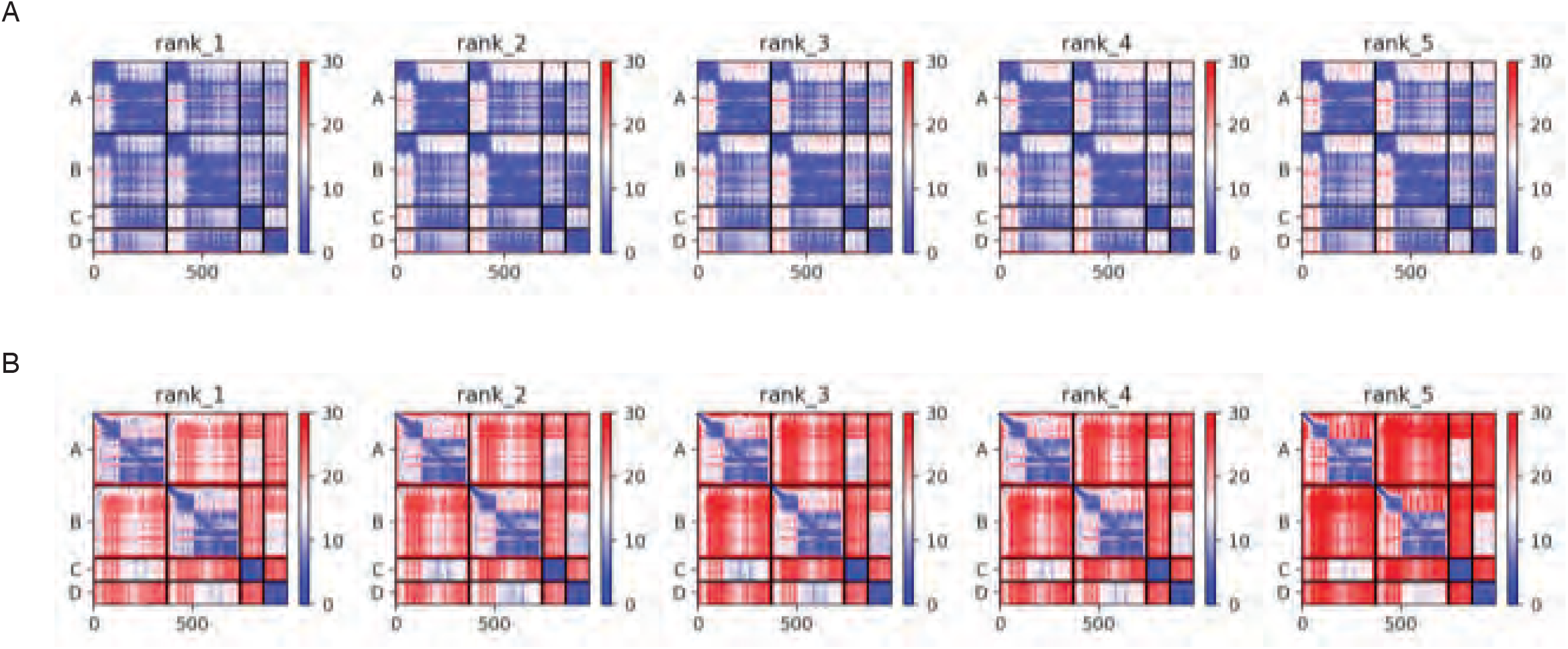
PAE plots of AlphaFold2 structure predictions. Predicted aligned error of the AlphaFold2 structure prediction of the following proteins: **A.** *B. subtilis* RsbU (chain A and B), and RsbV (chain C and D **B.** *B. subtilis* SpoIIE (chain A and B) and SpoIIAA (chain C and D).

**Supplemental Figure 8:**
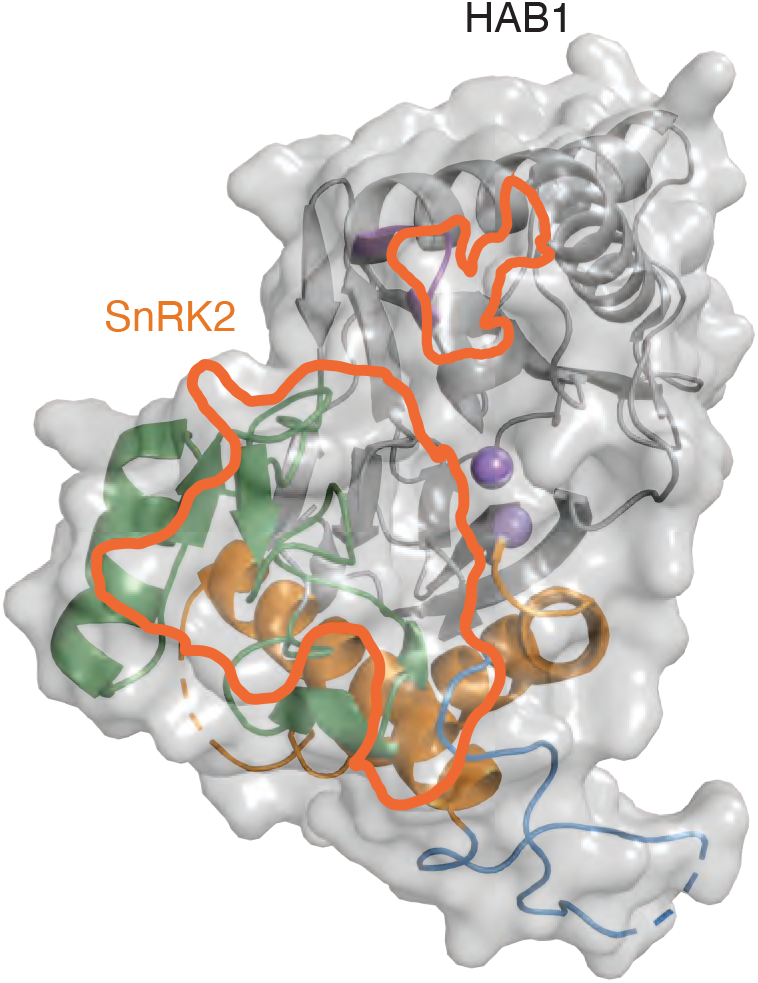
HAB1/SnRK2 interface show contact with flap, switch and α2/β4 loop. Crystal structure of *Arabidopsis thaliana* HAB1 depicting binding interface with SnRK2 outlined in orange based on a 1.4 Å probe radius. Interface outline shows SnRK2 makes contact with flap (green), switch (orange) and α2/β4 loop (blue).

